# A Cas9-fusion proximity-based approach generates an *Irak1-Mecp2* tandem duplication mouse model for the study of MeCP2 duplication syndrome

**DOI:** 10.1101/2023.02.07.527511

**Authors:** Eleonora Maino, Ori Scott, Samar Z. Rizvi, Shagana Visuvanathan, Youssif Ben Zablah, Hongbin Li, Ameet S. Sengar, Michael W. Salter, Zhengping Jia, Janet Rossant, Ronald D. Cohn, Bin Gu, Evgueni A. Ivakine

**Author notes:** These authors jointly supervised this work (Bin Gu, Evgueni A. Ivakine).

## Abstract

MECP2 duplication syndrome (MDS) is a neurodevelopmental disorder caused by tandem duplication of the *MECP2* locus and its surrounding genes, including *IRAK1*. Current MDS mouse models involve transgenic expression of *MECP2* only, limiting their applicability to the study of the disease. Herein, we show that an efficient and precise CRISPR/Cas9 fusion proximity-based approach can be utilized to generate an *Irak1-Mecp2* tandem duplication mouse model. The *Mecp2 Dup* model displays a neurological phenotype in keeping with MDS and demonstrates an abnormal immune response to infection not previously observed in other mouse models, possibly stemming from concurrent *Irak1* overexpression. The *Mecp2 Dup* mouse line thus provides an innovative tool to investigate disease mechanisms and potential therapeutic development.

## Introduction

MeCP2 is a transcriptional regulator with a multi-faceted role in the development and function of various tissues, most notably within the central nervous system (D’Mello, 2021). Mutations in *MECP2* underlie two pediatric neurodevelopmental disorders: Rett syndrome (RTT) caused by *MECP2* loss-of-function mutations (Amir et al., 1999), and *MECP2* duplication syndrome (MDS) resulting from duplications of the X chromosome region X28q (Van Esch et al., 2005). While the two disorders are caused by seemingly opposite genetic alterations, their phenotypes surprisingly overlap within the realm of neurodevelopmental disability, suggestive of a precise dosage requirement of MeCP2 within the nervous system. RTT affects 1:10,000 females and manifests as developmental regression, breathing abnormalities, seizures, anxiety, and metabolic dysfunction (Kyle et al., 2018). MDS affects mainly boys, with an estimated live birth prevalence of 1:150,000 (Giudice-Nairn et al., 2019), and is suspected to underly 1% of undiagnosed X-linked intellectual disabilities (Lugtenberg et al., 2009). It is characterized by developmental and psychomotor delay, locomotor and speech complications, autism spectrum disorder behaviour, epilepsy and seizure development, and susceptibility to infections (Ta et al., 2022). Life expectancy is drastically reduced, with patients often succumbing to respiratory tract infections (Friez et al., 2006).

From a genomic perspective, while RTT is primarily caused by point mutations affecting a single gene, MDS presents additional complexity as it can be caused by duplications of different sizes ranging from 50 kb to 15 Mb, affecting at least 2 genes: *IRAK1* and *MECP2* (Peters et al., 2019). As a result, faithful modeling of MDS remains a challenge. To date, MDS has primarily been investigated utilizing transgenic mice overexpressing the murine or human *MECP2* transgene, such as the *Tau-Mecp2* (Koerner et al., 2018; Luikenhuis et al., 2004), *MECP2-TG* (Collins et al., 2004) and hDup (Shao et al., 2021) mouse models. These models have been instrumental in elucidating certain pathophysiological mechanisms underlying the disease and demonstrating its reversibility (Sztainberg et al., 2015). However, none of these models recapitulate the genomic rearrangements harboured by most patients, caused by head-to-tail tandem duplication mutations. More importantly, they do not account for the duplication of *IRAK1*, which is shared by all MDS patients and which may influence disease manifestations through its critical role in host immunity (Gottipati et al., 2008).

The need to model genomic copy number variations (CNV) is not unique to MDS: indeed deletions and duplications of genomic regions underlie many human genetic disorders (Lupski, 2009). In particular, multi-genic tandem duplications are associated with several disorders causing intellectual disability, autism spectrum disorder and developmental delay (Coe et al., 2019; Takumi and Tamada, 2018; Zarrei et al., 2019). With the advent of new CRISPR/Cas9 genome editing technologies, mouse models recapitulating disease-causing CNVs have been generated. Current approaches have been shown to be quite efficient and precise in generating deletions. However, generating model organisms recapitulating tandem duplications is still an inefficient, inaccurate, and time-consuming process, and only a few such models have been generated (Maino et al., 2021; Pristyazhnyuk et al., 2019).

Here, we optimized the tandem duplication generation approach by utilizing a Cas9 fusion proximity-based strategy. We employed this methodology to generate the *Mecp2 Dup* mouse model, harbouring a 160 Kb tandem duplication on the X chromosome. This duplication region encompasses both the *Irak1* and *Mecp2* loci, known to be the minimal duplicated region shared by MDS patients. We showed that the *Mecp2 Dup* mouse model recapitulates the main neurobehavioral aspects of MDS and has an abnormal immune response to infection not previously described in the *Mecp2* overexpression models, possibly due to the impact of *Irak1* duplication. We conclude that the *Mecp2 Dup* mouse model is an excellent platform to better investigate disease mechanisms and develop novel therapeutic approaches for MDS patients. Furthermore, the fusion Cas9 proximity-based approach can be used more broadly as an efficient and precise method of generating models of complex genomic rearrangements.

## Materials and Methods

### Construct generation

For constructing the PCS2+ spCas9-FKBP plasmid, the FKBP coding sequence was amplified from the PM-FRB-Cerulean-T2A-FKBP-5-ptase plasmid (Addgene 40897, a kind gift from Dr. Peter Varnai) (Tóth et al., 2012) and used to replace the monomeric streptavidin (mSA) coding sequence of the PCS2+ Cas9-mSA plasmid (Addgene 103882) (Gu et al., 2018) using infusion cloning (Takara). For constructing the PCS2+ saCas9-FRB plasmid, the saCas9 and FRB coding sequences were amplified from the pX601 plasmid (Addgene 61591, a kind gift from Dr. Feng Zhang) (Ran et al., 2015) and the PM-FRB-Cerulean-T2A-FKBP-5-ptase plasmid (Addgene 40897, kind gift from Dr. Peter Varnai) (Tóth et al., 2012), respectively, and assembled as a c-terminal FRB fusion gene and replacing the spCas9-mSA cassette of the PCS2+ Cas9-mSA plasmid (Addgene 103882) (Gu et al., 2018) using infusion cloning (Takara).

### Producing mRNAs and sgRNAs for microinjection

mRNAs and sgRNAs were produced using in vitro transcription following our published protocol (Gu et al., 2020b). Briefly, to produce the SpCas9-FKB and the SaCas9-FRB mRNA for the microinjection, pCS2+ plasmids were linearized with NotI restriction digestion and used as template for in vitro transcription using the mMESSAGE mMACHINE^®^ SP6 Transcription Kit (Thermo Fisher Scientific). The sgRNAs were produced by utilizing the MEGAshortscript™ T7 Transcription Kit (Thermo Fisher Scientific). The RNeasy Mini Kit (Qiagen) was utilized to purify all RNA products.

### Producing bridge donor

A bridge donor was designed consisting of 136 bp upstream sequence and 186 downstream sequence from the putative junction sequence for the tandem duplication. An EcoRI restriction site is placed between the upstream and downstream sequences. The sequence of the bridging donor (Table S7) was synthesized as a gBlock from IDT. PCR products were prepared with a pair of primers (Table S8) and purified for microinjection following our published protocols (Gu et al., 2020b).

### Mouse model generation

The *Mecp2 Dup* and *Mecp2 Del* mouse models were generated by microinjecting two-cell stage embryos of the CD1 mouse strain following our published protocols (Gu et al., 2020b, 2020a). Microinjections were performed using a Leica microscope and micromanipulators (Leica Microsystem Inc.). Injection pressure was provided by a FemtoJet (Eppendorf) and negative capacitance was generated using a Cyto721 intracellular amplifier (World Precision Instrument). Microinjections were performed in M2 medium (Zenith Biotech). A mixture of SpCas9-FKB mRNA (100 ng/μl), SaCas9-FRB mRNA (100 ng/μl), *Irak1* sgRNA (50 ng/μl, protospacer sequence: TAGCATCAATCAGCCCTAGT (SpCas9 sgRNA)), *Tex28* sgRNA (50 ng/μl, protospacer sequence: CAGCTGTACTATGTTACCCAG, (SaCas9 sgRNA)), and bridging donor (5 ng/μl) was microinjected in nuclease-free injection buffer (10mM Tris-HCl, pH 7.4, 0.25mM EDTA). Embryos were cultured in 5nM rapamycin in KSOM (Zenith Biotech) for 6 hours and then implanted into the oviducts of E0.5 pseudo-pregnant females (30 embryos/female).

### Animal studies

All the animals utilized in this study were maintained in the specific-pathogen free facility at The Centre for Phenogenomics (TCP), Toronto, on a 12 h light/dark cycle and provided with food and water *ad libitum* in individually ventilated units (Techniplast). Only male mice on a CD-1 background aged from 6 up to 52 weeks of age were utilized in this study. Mice were randomly assigned to either the experimental or control group. Blind experimenters carried out the behavioural study described in the manuscript. All procedures involving animals were performed in compliance with the Animals for Research Act of Ontario and the Guidelines of the Canadian Council on Animal Care. The Institutional Animal Care Committee reviewed and approved all procedures conducted on animals at TCP.

### Genomic DNA isolation, PCR and RT-PCR

Genomic DNA was isolated using the DNeasy blood and tissue kit (Qiagen) according to the manufacturer’s protocol. PCR was performed using DreamTaq polymerase (Thermo Fisher Scientific) The primers utilized for PCR amplification are reported in Table S8.

### Sanger Sequencing

Amplified DNA was PCR-purified using QIAquick PCR Purification Kit (Qiagen) according to the manufacturer’s protocol and prepared for Sanger sequencing using the BigDye™ Terminator v3.1 Cycle Sequencing Kit (Thermo Fisher Scientific). Samples were sequenced on an Applied Biosystems SeqStudio Genetic Analyzer (Thermo Fisher Scientific), and electropherograms were analyzed using SnapGene software.

### RNA isolation, RT-PCR and quantitative PCR

RNA extraction was performed using Trizol reagent (Thermo Fisher Scientific) following the manufacturer’s protocol. Next, 1 μg of mRNA was reverse transcribed using SuperScript III reverse transcriptase kit (Thermo Fisher Scientific). Quantitative PCR was performed using the fast SYBR Green master mix (Qiagen) on a StepOnePlus real-time PCR (Applied Biosystems). ΔΔCt was analyzed to assess fold changes between mutant and wild type mice.

### Protein isolation and western blot

Mouse tissue was homogenized in 400 μl of RIPA homogenizing buffer (50-mM Tris HCl pH 7.4, 150-nM NaCl, 1-mM EDTA, supplemented with protease-inhibitor cocktails (Roche)) and lysed with a MagNA Lyser. Subsequently, 400 μl of RIPA double-detergents buffer (2% deoxycholate, 2% NP40, 2% Triton X-100 in RIPA homogenizing buffer) was added to the lysates, which were then incubated for 45 min at 4 °C, and then centrifuged for 10 min at 13,000 rpm. Protein concentration was measured using a BCA Assay (Thermo Fisher Scientific). Protein was separated on a 4–12% Bis-Tris gel and transferred using iBlot 2 transfer apparatus (Thermo Fisher Scientific). A 5% milk solution in TBST was used for blocking for 1 hour at room temperature. The membrane was then incubated with appropriate primary antibodies: rabbit recombinant anti-MECP2 (ab253197, Abcam, 1:3000), rabbit monoclonal D51G7 anti-IRAK1 (#4504, Cell Signaling, 1:1000), rabbit monoclonal anti-GAPDH (sc-47724, Santa Cruz, 1:10000) overnight at 4C. The membranes were then incubated 1 hour at room temperature with horseradish peroxidase conjugated goat anti-rabbit IgG (Abcam, ab6721). Signal detection was achieved using SuperSignal West Femto Maximum Sensitivity Substrate (Thermo Fisher Scientific) according to the manufacturer’s protocol.

### Digital droplet PCR (ddPCR)

Genomic DNA extracted from mouse tails was utilized to perform ddPCR by The Centre for Applied Genomics (TCAG) at the Hospital for Sick Children. Copy number estimation of *Tex28* and *Mecp2* were performed using the QX200 Droplet Digital PCR system (Bio-Rad Laboratories, Inc.) using Taqman Copy Number assays; Mm00631912_cn and Mm00629942_cn (Life Technologies). Prior to the copy number experiment, 100 ng of genomic DNA was digested with 5U of BtsCI in a 3 μl reaction (New England Biolabs), 1 hour at 50°C incubation and no enzyme denaturation. The 20 μl copy number reaction mix consisted of 10 μl of 2x ddPCR SuperMix for Probes (Bio-Rad Laboratories), 1 μl of the Copy Number target assay (labeled with FAM), 1ul of the Copy Number Reference assay (Mouse *Tfrc*, Life Technologies, part 4458366, labeled with VIC), 5 μl water and 3 μl of the digested genomic DNA. The Copy Number assay was validated by temperature gradient to ensure optimal cluster separation of target and reference droplets. Cycling conditions for the reaction were 95C for 10 min, followed by 45 cycles of 94C for 30 sec and 60C for 1 min, 98C for 10 minutes on a Life Technologies Veriti thermal cycler. Data was analyzed using QuantaSoft v1.4 (Bio-Rad Laboratories). Wildtype controls and non-template controls were included with each run.

### Whole Genome Sequencing

DNA extracted from mouse tails was utilized for whole genome sequencing (WGS), which was performed using the Illumina HiSeq X system (San Diego, CA, USA) by TCAG (Hospital for Sick Children). In brief, 400 ng of DNA sample was used for library preparation using the Illumina TruSeq PCR-free DNA Library Prep Kit, where DNA was sonicated into an average of 350-bp fragments. A-tailed and indexed TruSeq Illumina adapters were ligated to end-repaired sheared DNA fragments before the library was amplified. Libraries were analyzed using Bioanalyzer DNA High Sensitivity chips (Agilent Technologies, Santa Clara, CA, USA) and quantified using qPCR. The libraries were loaded in equimolar quantities and pair-end sequenced on the Illumina HiSeq X platform to generate 150-bp reads. Integrative Genomics Viewer (IGV) version 2.8.2 was used for analysis with GRCm38/mm10 as the murine reference genome.

### RNA Sequencing

RNA was extracted from mice hippocampi and quantified using a Qubit RNA HS assay (Thermo Fisher Scientific). RNA sequencing was performed by the TCAG (The Hospital for Sick Children) using the Illumina HiSeq 2500 system, producing 150-bp paired-end reads. Briefly, raw transcript reads were aligned to the GRCm38/mm10 mouse reference genome using STAR aligner, v.2.6.0c. (https://github.com/alexdobin/STAR). HTSeq v.0.6.1p2 was used to determine the absolute number of read counts for each gene. Normalization and differential expression analysis were completed using the R package DESeq2 v.1.26.0s package (https://bioconductor.org/packages/release/bioc/html/DESeq2.html). Initial minimal filtering of 10 reads per gene for all samples was applied to the datasets. False discovery and multiple testing were controlled for, and an adjusted *P* value was computed using the Benjamini–Hochberg method. Differentially expressed genes (DEGs) were defined as genes with base mean >10, and false discovery rate less than 15%. DEGs list and fold change are reported in Table S2. Hierarchical clustering was computed using Euclidean distance as distance measurement method, and average linkage as clustering method. Gene ontology analysis of the DEGs was performed using g:Profiler (https://biit.cs.ut.ee/gprofiler/gost). The complete gene ontology list is reported in Table S3.

### Open field

For the open field test, mice were placed in the frontal center of a transparent Plexiglas open field (41.25 cm × 41.25 cm × 31.25 cm) illuminated by 200 lx. The VersaMax Animal Activity Monitoring System recorded activity in the center and periphery of the open field arena for 20 min per animal.

### Rotarod test

The mice were placed on a rotating rod (Panlab) that accelerated from 4 to 40 revolutions per minute. The duration of each trial was a maximum of 300 seconds for the *Mecp2 Dup* mice, and a maximum of 600 seconds for the *Mecp2 Del* mice. Mice were tested for 3 consecutive days, 3 trials each, with an interval of 15 min between trials to rest. The time that it took for each mouse to fall from the rod (latency to fall) was recorded.

### Long Term Potentiation measurements

We anesthetized 6-weeks-old mice with urethane (20% wt/vol, intraperitoneal (i.p.)). We prepared parasagittal hippocampal slices (300 μm) in ice-cold ACSF and placed them in a holding chamber (30°C) for 40 min, and then allowed it to passively cool down to room temperature (21 to 22°C for ≥ 30 min) before recording. We transferred a single slice to a recording chamber and perfused with artificial cerebrospinal fluid (ACSF) (Sigma-Aldrich) at 4 ml/min composed of 124mM NaCl, 2.5mM KCl, 1.25mM NaH_2_PO_4_, 2mM MgCl_2_, 11mM D-glucose, 26mM NaHCO_3_ and 2mM CaCl_2_ saturated with 95% O_2_ (balance 5% CO_2_) at room temperature (pH 7.40, 305 mOsm). We evoked synaptic responses by stimulating Schaffer collateral afferents using bipolar tungsten electrodes located ~50 μm from the pyramidal cell body layer in CA1. We recorded extracellular fEPSPs using ACSF-filled glass micropipettes placed in the stratum radiatum 60–80 μm from the cell body layer. Testing stimuli (0.1 ms in duration) were delivered at a frequency of 0.033 Hz to evoke a half-maximal fEPSP. In LTP experiments, theta-burst stimulation (TBS) consisted of 16 bursts of four pulses at 100 Hz, delivered to Schaffer collateral afferents at an inter-burst interval of 200 ms. We amplified raw data using a MultiClamp 700B amplifier and a Digidata 1322A acquisition system sampled at 10 kHz, and analyzed the data with Clampfit 10.6 (Axon Instruments) and Sigmaplot 11 software.

### Fear conditioning test

Individual pre-handled mice were placed in a conditioning chamber with controlled contextual cues and an electrified shock floor (Omnitech Electronics Inc., Columbus, Ohio, USA). Each individual mouse was allowed to explore the conditioning chamber for 3 minutes before a conditioned stimulus, a 30-second tone (95 dB, 3600 Hz) was played for 30 seconds. During the last 2 seconds of the conditioned stimulus duration, an unconditioned stimulus (a mild foot shock of 0.75 mA) was delivered. Five training sessions were delivered in succession with an interval period of 30 seconds between the training sessions. After the last training session, the mouse was allowed to remain in the chamber for 30 seconds to assess the behavior of the mouse after the training. Fear responses were assessed by recording the freezing behaviors of the mouse using FreezeView2 (Coulbourn Instrument, Whitehall, Pennsylvania, USA). Freezing was defined as lack of any locomotor activity except for slight head and tail movements (all within an index of motion of 10). Two hours and 24 hours following the training, the fear memory of the mouse was tested using contextual and cued tests. During the contextual test, the contextual cues that were present during conditioning remained. The mouse was placed in the recording chamber for 4 minutes. During the cued test, the chamber and the contextual cues were changed. Following a two minutes acclimation, the same tone was played for 2 minutes. The mouse’s movement was tracked during both contextual and cued test, and freezing responses were measured using FreezeView2. The duration of freezing responses of the mouse was measured and compared to the total duration of the test. Freezing responses were presented as a percentage.

### Influenza virus administration

*Mecp2 Dup* mice and wildtype littermates at 10 weeks of age received intranasal infection with Influenza H1N1 virus at a dosage of 100pfu/35g mouse in PBS. Mice were anesthetized via intraperitoneal injection of ketamine/xylazine prior to infection. The dose utilized was previously determined at LD50 for CD-1 wildtype mice.

### Blood collection

The blood was collected by saphenous vein puncture, cheek bleed or cardiac puncture (for endpoint measurements).

### Blood cells count

Blood cell count analysis at baseline levels was performed by TCP with the Hemavet Multispecies Hematology Systems - HV950 (Drew Scientific, Inc.). Blood cell count measurements at day 4 post infection were performed by Biovet. EDTA infused whole blood at room temperature were used for the analysis. The analysis was performed on the day of the collection.

### Bronchoalveolar fluid (BALF) collection

After mice were euthanized, a small incision made in the trachea, a needle wrapped with soft plastic tubing was inserted into the trachea, secured with sutures, and used to administer 0.5 ml PBS to lavage the lungs. The collected supernatants were used for cytokines analysis.

### Chemokines and cytokines analysis

BALF (undiluted) and serum samples (1:2 dilution in PBS) were collected and stored at −80°C until analysis. Samples were submitted to and analyzed utilizing the Mouse Multiplex Cytokine Array / Chemokine Array 44-Plex (MD44) by Eve Technologies.

### Statistical analysis

All statistical analyses were performed using GraphPad Prism (GraphPad software). Data normality was tested using Shapiro-Wilk test for n<8 datasets and D’Agostino-Pearson test for n>8 datasets. Parametric data were analyzed with two-tailed Student’s *t*-tests, one-way ANOVA followed by Tukey’s post hoc test, or two-way ANOVA repeated measures followed by

Bonferroni multiple comparisons correction. Datasets with non-normal distribution were analyzed using two-tailed Mann-Whitney U tests or Kruskal–Wallis test followed by Dunn’s post hoc test. The exact numbers of animals used, and the specific statistical analysis utilized are indicated in each figure legend. All behavioural analyses were performed blinded to genotype.

## Results

### A fusion Cas9 proximity-based approach generates the *Mecp2 Dup* and *Mecp2 Del* mouse models

It is essential to optimize current strategies to generate mouse models recapitulating disease-causing structural genomic variants, including large deletions and in particular, tandem duplications. We hypothesized that the efficiency of CNV generation would be enhanced by promoting the physical proximity of the DNA ends involved in the rearrangement. The approach we developed utilizes the ability of the FKBP and FRB domains to dimerize upon rapamycin binding (Banaszynski et al., 2005). Our system consists of two orthogonal Cas9 enzymes targeting the 3’ and 5’ ends of the region of interest (Fig 1A). A *Staphylococcus aureus* (SaCas9) is fused with the FK506 binding protein-12 (FKBP) (SaCas9-FK506), paired with a single-guide RNA (sgRNA) targeting the 3’ region of the desired rearrangement. Concurrently, *Streptococcus pyogenes* Cas9 (SpCas9) is fused with the FKBP-rapamycin-binding (FRB) domain of mTOR (SpCas9-FRB), with a sgRNA targeting the 5’ region. Upon rapamycin treatment, the FKBP and FRB domains dimerize bringing the two Cas9 and the free DNA ends in close proximity, and promoting the occurrence of the desired CNV. To favour the formation of a precise duplication junction, a double-stranded DNA PCR product spanning the duplication junction was added to the system. Furthermore, we reasoned that performing the above-described genetic modification during the long G2 phase of the cell cycle of 2-cell stage mouse embryos would improve the efficiency of tandem duplication, due to the presence of closely aligned sister chromatids (Gu et al., 2018).

**Figure 1.**
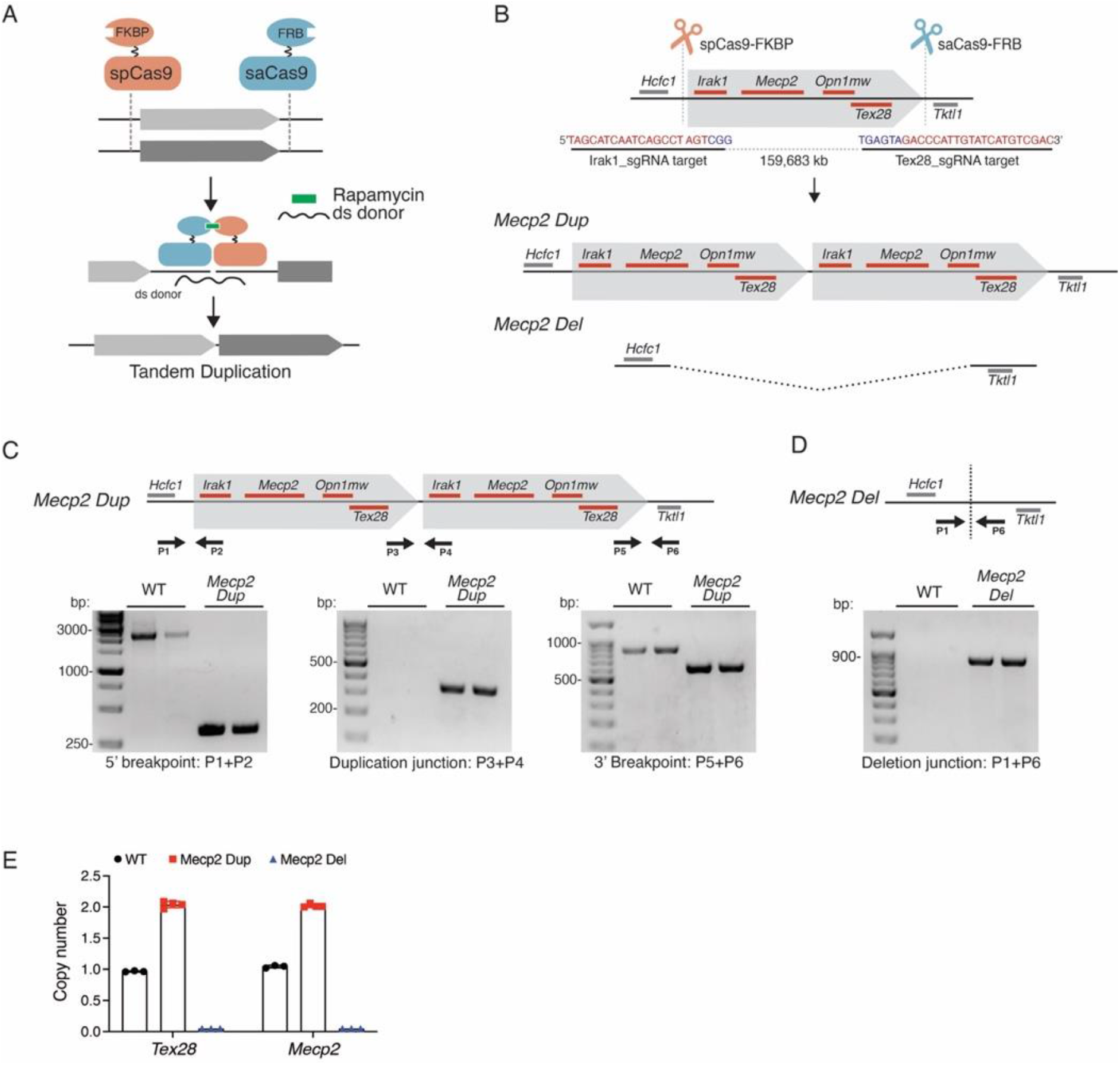
A fusion Cas9 proximity-based approach generates the *Mecp2 Dup* and *Mecp2 Del* mouse models. **A)** Schematics of the proximity-based approach to generate tandem duplications. Upon rapamycin treatment, the two Cas9 dimerize bringing in proximity the DNA ends to rearrange. A double-stranded donor (ds donor) bridging the duplication junction is micro-injected together with the CRISPR/Cas9 components. **B)** A ~160 kb region encompassing the genes *Irak1, Mecp2, Opn1mw* and *Tex28* was targeted to generate the *Mecp2 Dup* and *Mecp2 Del* mouse models. The target sequences for both guides utilized are shown, with the PAM sequences denoted in blue and the guide sequence in red. Cas9 and the sgRNAs are represented as scissors. **C)** PCR amplification of the duplication 5’ and 3’ breakpoints and junction in the *Mecp2 Dup* founder mouse to confirm WGS. Primers are represented by arrows. **D)** PCR amplification deletion junction in the *Mecp2 Dup* founder mouse. Primers are represented by arrows. **E)** ddPCR confirmed the copy number of the duplicated or deleted region with probes targeting *Tex28* and *Mecp2*. A probe targeting the *Tfrc* gene was utilized as reference.

We applied this methodology to generate a mouse model recapitulating a tandem duplication mimicking MD patients’ causing mutations. We designed two sgRNAs, referred to as Irak1_sgRNA and Tex28_sgRNA, flanking the ~160 kb target region for duplication, which includes the genes *Irak1, Mecp2, Opnmw1* and *Tex28* (chrX:74,012,497-74,172,198) (Fig 1B). We microinjected Cas9 mRNAs, sgRNAs and the 331 bp bridging donor into 180 2-cell stage embryos, following the previously established 2C-HR-CRISPR protocol (Gu et al., 2018). Embryos were treated with rapamycin for 6 hours to allow dimerization, and then transferred to pseudo-pregnant females. Twenty-four live born pups were produced. The pups were screened using PCR amplifying the putative tandem duplication junction. Since a two-guide approach is a well-established technique to generate genomic deletions, we screened the newborn mice also for the presence of the reciprocal *Irak1-Tex28* deletion. The tandem duplication and deletion generation efficiency observed were 3/24 (12%) and 16/24 (66%), respectively. One mouse contained both a tandem duplication and a large deletion, suggesting reciprocal *Irak1-Tex28* duplication/deletion. This founder mouse was outcrossed to wildtype mice to establish the *Mecp2 Dup* and *Mecp2 Del* mouse lines, respectively harbouring a *Irak1-Tex28* duplication and deletion.

Given that CRISPR/Cas9 may generate mosaic mutations, we conducted comprehensive genetic quality control in N1 progenies as described previously (Gu et al., 2022). Because inadvertent structural variations may be generated by the editing process, the *Mecp2 Dup* N1s were analyzed by whole genome sequencing (WGS), which confirmed the presence of the intact tandem duplication. Minimal rearrangements were identified at both the duplication junction and breakpoints, which represent the guide target sites (Table S1). PCR followed by Sanger sequencing confirmed a 2026 bp deletion around the Irak1_sgRNA target site and a 252 bp deletion at the Tex28_sgRNA target site (Fig 1C, Fig S1B-C). The duplication junction harbours a 170 bp insertion compared with the predicted junction (Fig S1A). These minor insertions/deletions did not impact the expression of the primary MDS-defining genes *Mecp2* or *Irak1*. Furthermore, genetic changes in MDS patients represent non-recurrent duplications with inconsistent junction sequences, thus these minor changes are unlikely to affect the phenotypes of our mouse model. PCR followed by Sanger sequencing was utilized to validate the presence of the predicted deletion junction in the *Mecp2 Del* mouse model (Fig 1D, Fig S1D). Digital droplet PCR (ddPCR) confirmed the predicted copy number of the region in *Mecp2 Dup, Mecp2 Del* and wildtype control mice (Fig 1E).

The deletion of *Mecp2* resembles the MECP2 loss of function mutations seen in the context of RTT. Accordingly, *Mecp2 Del* mice develop a phenotype compatible with RTT including a reduced lifespan, compromised motor coordination, and clasping behaviour (Fig S2A-D). In summary, our Cas9 fusion proximity-bases approach generated the first tandem duplication mouse model of MDS, coined the *Mecp2 Dup* mouse, as well as an RTT mouse model known as the *Mecp2 Del* mouse, which will be used as a control for our experiments.

### The *Mecp2 Dup* mouse model exhibits increased *Mecp2* and *Irak1* expression

Molecular characterization of the *Mecp2 Dup* mice began with confirming the expression of both *Mecp2-e1* and *Mecp2-e2* isoforms in the brains of the *Mecp2 Dup* mice (Fig 2A-B). To evaluate the levels of *Mecp2* expression, we performed qRT-PCR in various brain areas, including hippocampus, cortex, and cerebellum, identifying a 2-2.7 fold-increased expression compared to wildtype littermates across all of the regions analysed (Fig 2C, Fig S3A). *Irak1* expression was 2-2.3-fold-increased in the brain of *Mecp2 Dup* compared with wildtype controls (Fig 2D, Fig S3B). Transcript analysis confirmed the absence of both *Mecp2* and *Irak1* expression in the brain of *Mecp2 Del* mice (Fig 2C-D). MECP2 and IRAK1 protein levels were evaluated in the aforementioned mentioned brain areas via immunoblotting with values correlating with the transcript levels for both MECP2 and IRAK1, ranging from 2.3-3.7-fold and 1.8-2.7-fold compared to wildtype, respectively (Fig 2D-E, Fig S3E-H).

**Figure 2.**
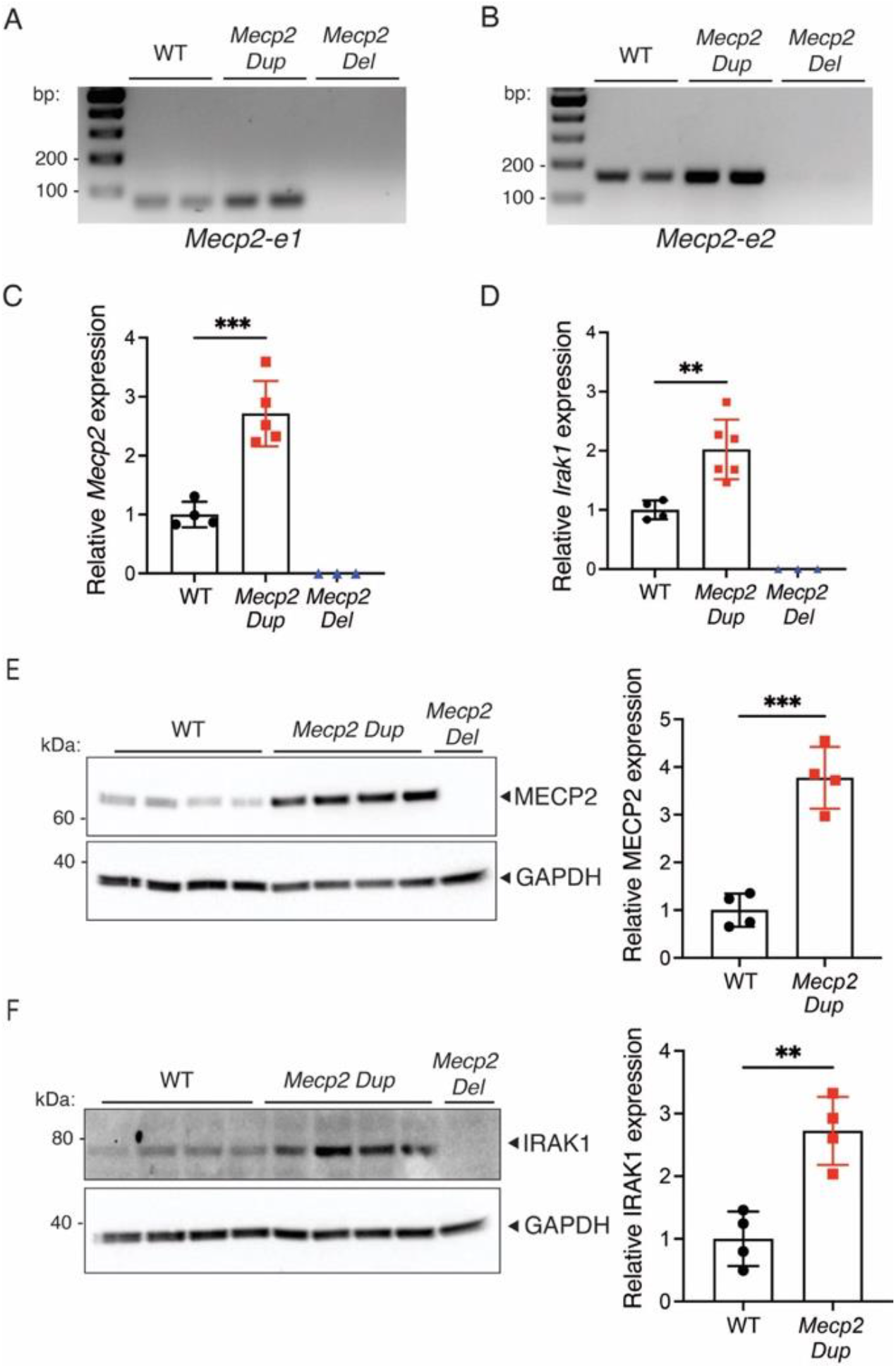
*Mecp2* and *Irak1* are overexpressed in the hippocampus of *Mecp2 Dup* mice. RT-PCR analysis performed in the hippocampus detected the expression **A)** *Mecp2-e1* and **B)** *Mecp2-e2* expression in the hippocampus of wild type and *Mecp2 Dup*. No *Mecp2* was observed in *Mecp2 Del* mice hippocampi. **C)** The levels of *Mecp2* expression were analyzed via qPCR in 10 weeks *Mecp2 Dup* mice, wild type littermates and *Mecp2 Del* mice. The data is normalized over *Gapdh* expression. WT, n=4; *Mecp2 Dup*, n=5; *Mecp2 Del*, n=3. **D)** *Irak1* transcript levels were analyzed via qPCR. The data is normalized over *Gapdh* expression. WT, n=4; *Mecp2 Dup*, n=6; Mecp2 Del, n=3. Western blot analysis confirmed increased **E)** MECP2 and **F)** IRAK1 expression in the hippocampus of *Mecp2 Dup* mice compared to wild type littermates. GAPDH serves as a loading control (right). Densitometry analysis to quantify the amount of MECP2 and IRAK1 expression (left). WT, n=4; *Mecp2 Dup*, n=4; *Mecp2 Del*, n=1. All data are represented as the mean +/− SD. Statistical analyses were performed with Student’s t-test. **P<0.01, ***P<0.001.

### *Mecp2 Dup* mice develop a progressive neurological phenotype compatible with MDS

We evaluated the activity of *Mecp2 Dup* mice and wildtype littermates at 10, 20 and 52 weeks of age utilizing the open field test. Starting at 10 weeks, *Mecp2 Dup* mice showed increased activity for every parameter analyzed on the open field test, with symptoms progressing as the mice aged. At 10 weeks of age *Mecp2 Dup* mice showed a 20% increase in total distance travelled versus wild type mice (p<0.01), 60% increase in vertical activity (p<0.001) and 13% increase in average speed (p<0.05) in the open field arena (Fig 3A-C). At 20 weeks of age, these parameters increase up to 35% (p<0.0001), 84% (p<0.01) and 29% (p<0.0001), respectively, and remained stable up to 52 weeks of age (Fig 3A-C). In addition, the *Mecp2 Dup* mice spent up to twice the amount of time in the centre of the open field arena compared with wildtype littermates at every time point analyzed (Fig 3D). These data suggest that *Mecp2 Dup* mice display reduced anxiety compared to wildtype mice. In support of this observation, we also observed abnormal levels of activity in the centre of the open field arena, where MDS mice travelled 1.5-2.2 times more distance compared to wildtype mice, and showed 2.3-4.5 times more vertical activity (Fig S4A-B). *Mecp2 Dup* mice also spent 40-50% less time resting in the periphery of the open field arena (Fig S4C). We utilized a 3-day accelerated rotarod test to analyze motor coordination and cerebellar learning at 10, 20 and 52 weeks of age. Minimal differences were observed at 10 and 20 weeks of age (Fig S5A-B). However, at 52 weeks the *Mecp2 Dup* mice showed reduced motor coordination compared with wildtype littermates, falling from the rotarod 37-48% (p<0.5, p<0.01) earlier each day of the test (Fig 3E). No significant differences in body weight of MDS mice and wildtype controls were detected up to 38 weeks of age, when the *Mecp2 Dup* mice became significantly lighter (Fig S6A). Notably, *Mecp2 Dup* mice showed a 15-22% increase in brain-to-body weight ratios (p<0.01) at each timepoint analyzed (Fig 3E). The *Mecp2 Dup* mice had a reduced life span (Mantel-Cox test, p<0.01), with 40% mortality between 18 and 52 weeks of age, compared to wildtype littermates for which no deaths were observed. (Fig 3F).

**Figure 3.**
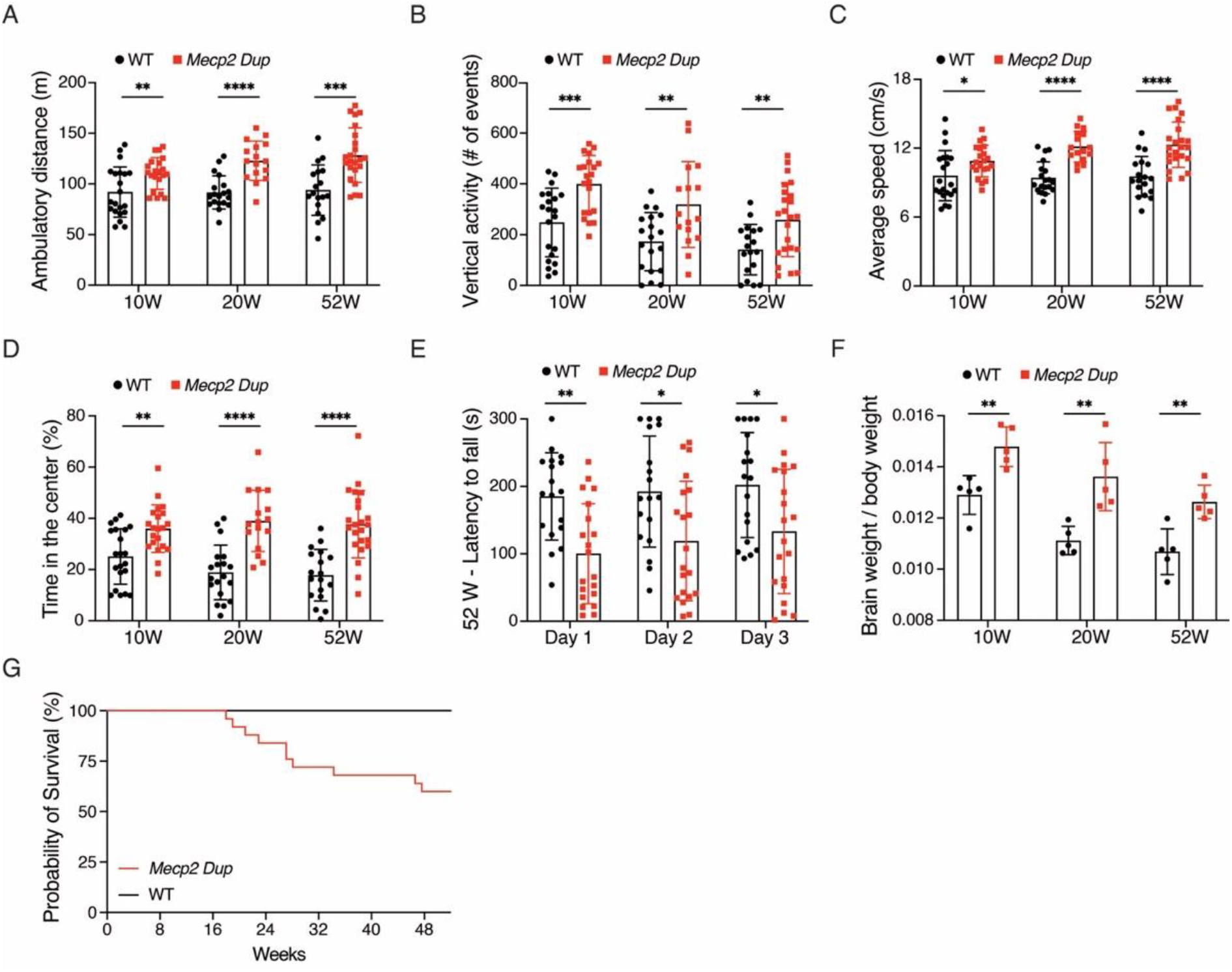
The *Mecp2 Dup* mice develop progressive neurological phenotypes. *Mecp2 Dup* mice showed increased activity and reduced anxiety-related behaviours on the open field test compared to wild type littermates at 10, 20 and 52 weeks of age. Parameters measured include **A)** Ambulatory distance, **B)** Vertical activity, **C)** Average speed, **D)** Time spent in the central area of the arena. *Mecp2 Dup*, n=16-23; WT, n=18-21. Statistical analyses were performed with Student’s t-test. **E)** A 3 days rotarod test identified reduced motor coordination in the *Mecp2 Dup* mice at 52 weeks of age. *Mecp2 Dup*, n=20; WT, n=18. Data was analyzed by two-way ANOVA repeated measures followed by Bonferroni’s multiple comparison test. **F)** Brain-to-body weight ratio was measured at every time point. *Mecp2 Dup*, n=5; WT, n=5. Statistical analyses were performed with Student’s t-test. **G)** Survival analysis of *Mecp2 Dup* mice up to 52 weeks of age was reduced compared to wild type littermates (** P=0.0084). *Mecp2 Dup*, n=23; WT, n=13. Survival curves were compared with the Mantel-Cox test. All data are represented as the mean +/− SD. *P<0.5, **P<0.01, ***P<0.001, **** P<0.0001.

### *Mecp2 Dup* mice show enhanced synaptic plasticity and abnormal learning

To evaluate hippocampal-related phenotypes in the *Mecp2 Dup* mice, we investigated neurotransmission at the Schaffer collateral synapses in the CA1 region of the hippocampus. To determine whether the overexpression of *Mecp2* might affect basal synaptic transmission, we recorded field excitatory postsynaptic potentials (fEPSPs) and found that the stimulation intensity-response relationship did not differ between *Mecp2 Dup* mice and wildtype littermates (Fig 4A). The *Mecp2 Dup* mice showed enhanced short-term synaptic plasticity compared to wildtype littermates as determined by paired-pulse facilitation (PPF) of the fEPSPs (Fig 4B).

**Figure 4.**
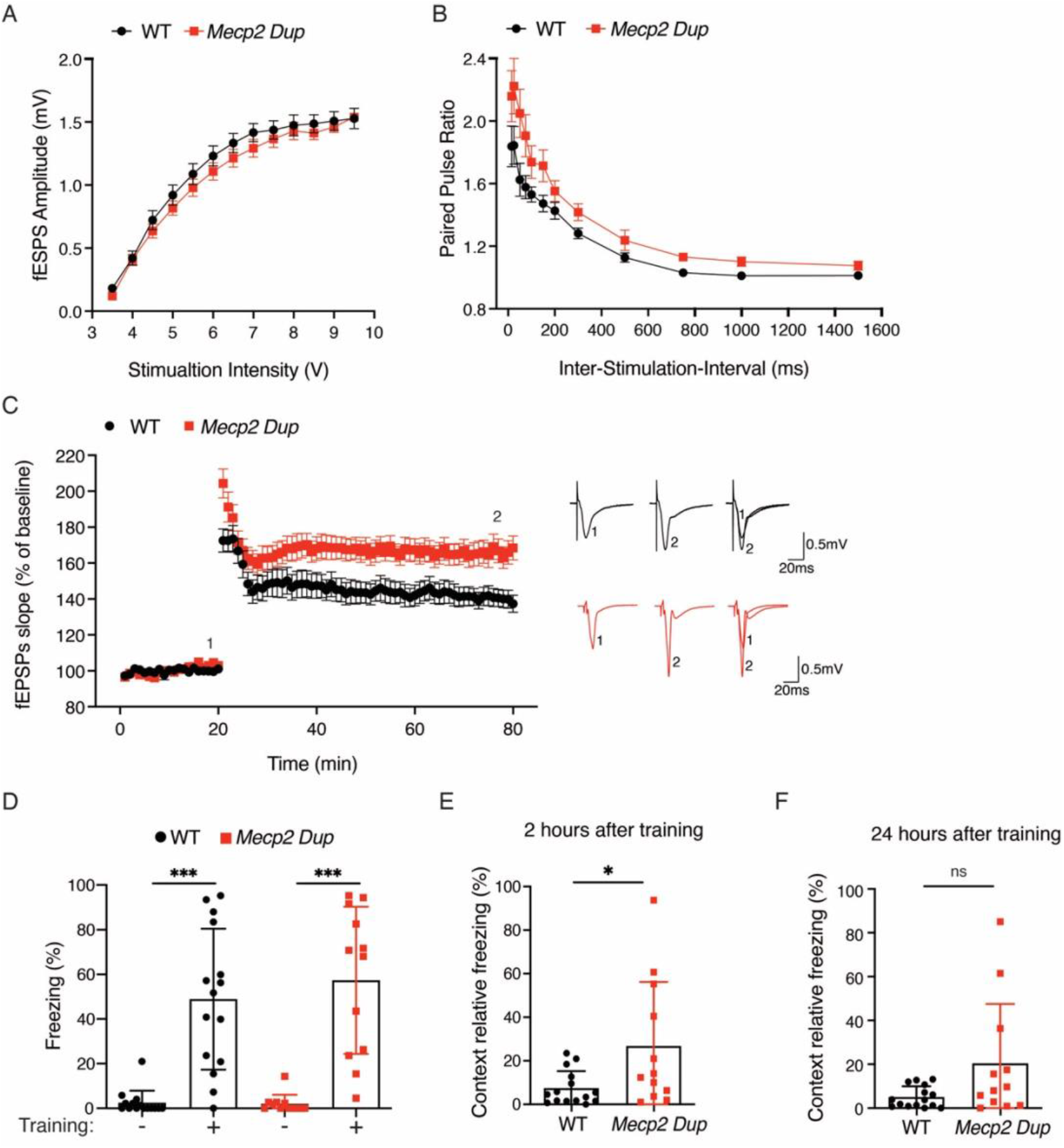
Electrophysiology analysis and fear conditioning test show abnormalities in hippocampal function in the *Mecp2 Dup* mice. **A**) Scatterplot of fEPSP peak amplitudes as a function of stimulus intensity for each genotype. *Mecp2 Dup*, n=17; WT, n=17. Data are represented as mean +/− SEM. **B)** Scatterplot of the paired pulse ratio as a function of the inter-burst interval for each genotype. *Mecp2 Dup*, n=11; WT, n=8. Data are represented as mean +/− SEM. *Mecp2 Dup* mice show increased paired pulse ratio at interstimulus intervals 15 ms (p<0.05), 25 ms (p<0.001), 50 ms (p<0.001), and 75 ms (p<0.001). Statistical analysis performed with two-way ANOVA followed by Sidak multiple comparisons test. **C)** Scatterplot of the normalized fEPSP slope over time induced by TBS of the Schaffer collateral inputs in hippocampal slices from each genotype. Traces from each genotype are represented as an average of fifteen consecutive fEPSPs recorded at the indicated times (time points 1 and 2). The magnitude of synaptic potentiation was as follows: WT (140.9 ± 4.9%, n = 17) and *Mecp2 Dup* (166.3 ± 6.5%, n = 15). *P = 0.004* as determined by Student t-test. Data values are mean +/− SEM. **B)** In conditioned fear analysis, there was no statistical difference in response to training at 20 weeks of age between *Mecp2 Dup* mice and wild type littermates. *Mecp2 Dup*, n=12; WT, n=15. Statistical analysis performed by Kruskal Wallis test followed by Dunn’s multiple comparison test. **C)** Percent freezing was enhanced in *Mecp2 Dup* compared to wild type when tested on contextual fear conditioning 2 hours after training. *Mecp2 Dup*, n=12; WT, n=15. Statistical analyses were performed with Student’s t-test. Data are represented as the mean +/− SD. *p<0.05. **D)** Contextual fear conditioning test performed 24 hours after training showed no statistical differences compared to wild type mice (p=0.1223). This suggests that mutant mice have enhanced short-term memory, while long-term memory is similar to wild type mice. *Mecp2 Dup*, n=12; WT, n=15. Statistical analyses were performed with Mann–Whitney *U* test. All data are represented as the mean +/− SD.

Moreover, we evaluated the capacity of Schaffer-collateral synapses to elicit long-term potentiation (LTP). *Mecp2 Dup* mice showed a robust LTP enhancement immediately after theta burst stimulation (TBS) that persisted for at least 60 min (Fig 4C) indicating increased synaptic plasticity compared to wildtype littermates. To determine the possible impact of these findings on hippocampal learning, we performed fear conditioning analysis in 20-week-old mice. No differences were observed in the freezing response during training between the two genotypes (Fig 4D). We then tested the mice 2 and 24 hours after training both with contextual and cued tests. The *Mecp2 Dup* mice showed increased contextual freezing presenting 3.6 (p<0.05) and 4 times (p = 0.1223) more freezing than wildtype at 2 and 24 hours, respectively (Fig 4E-F). Increased freezing behaviour, albeit not statistically significant, was noted on the cued test as well (Fig S7A-B).

### Alterations in *Mecp2* dosage lead to hippocampal transcriptional changes in *Mecp2 Dup* mice

As the *Mecp2 Dup* model develops hippocampal-related phenotypes and MeCP2 is a known global transcriptional regulator for hippocampal genes (Chahrour et al., 2008), we analyzed transcriptomic changes in the hippocampus via RNA-sequencing (RNA-seq), comparing *Mecp2 Dup*, *Mecp2 Del* and wildtype littermates at 6 weeks of age. Our analysis identified 339 differentially expressed genes in *Mecp2 Dup* versus *Mecp2 Del*, 59% of which were upregulated in *Mecp2 Dup* mice compared to *Mecp2 Del* mice (Fig 5A, Table S2). As expected, *Mecp2* and *Irak1* were identified as the top differentially-expressed genes (*Mecp2;* p=2.11e-145; *Irak1* p= 3.29e-104). Transcriptomic analysis identified several genes that overlapped with previous RNA-seq and microarray datasets performed in the context of *MECP2-related* disorders (Chahrour et al., 2008; Samaco et al., 2012; Sztainberg et al., 2015). Gene ontology analysis revealed that significantly enhanced pathways in *Mecp2 Dup* hippocampi are mainly including genes involved in transmembrane ion transport localized to synapses and neuronal projections. (Fig 5B, Table S3). We proceeded to validate some of top differentially-expressed genes with qRT-PCR (Fig 5C-F). Among these we demonstrated an increased expression in hippocampi of *Mecp2 Dup* mice compared to wild type littermates of transcripts for the Glutamate Decarboxylase 2 (*Gad2*) enzyme involved in γ-amino-butyric-acid (GABA) synthesis, previously shown to be regulated by MeCP2 (Chao et al., 2010); the Calcium Voltage-Gated Channel Subunit Alpha1 G (*Cacna1g*) associated with intellectual disability and epilepsy (Berecki et al., 2020; Chemin et al., 2018; Striessnig, 2021); the FXYD Domain Containing Ion Transport Regulator 7 (*Fxyd7*) which modulates the activity of the NA/K-ATPase (Meyer et al., 2020), consistently reported to be upregulated in the context of MDS and downregulated in RTT (Ben-Shachar et al., 2009); and decreased expression of the Poly(RC) Binding Protein 3 (*Pcbp3*), found to be dysregulated in patients with neurodevelopmental defects (Duijkers et al., 2019) and Down syndrome (Jagadeesh et al., 2020). Taken together, our results highlight how transcriptional changes in the hippocampus globally alter the expression of ion channel proteins, providing a molecular basis for the alteration in synaptic response that has been observed in MDS mouse models.

**Figure 5.**
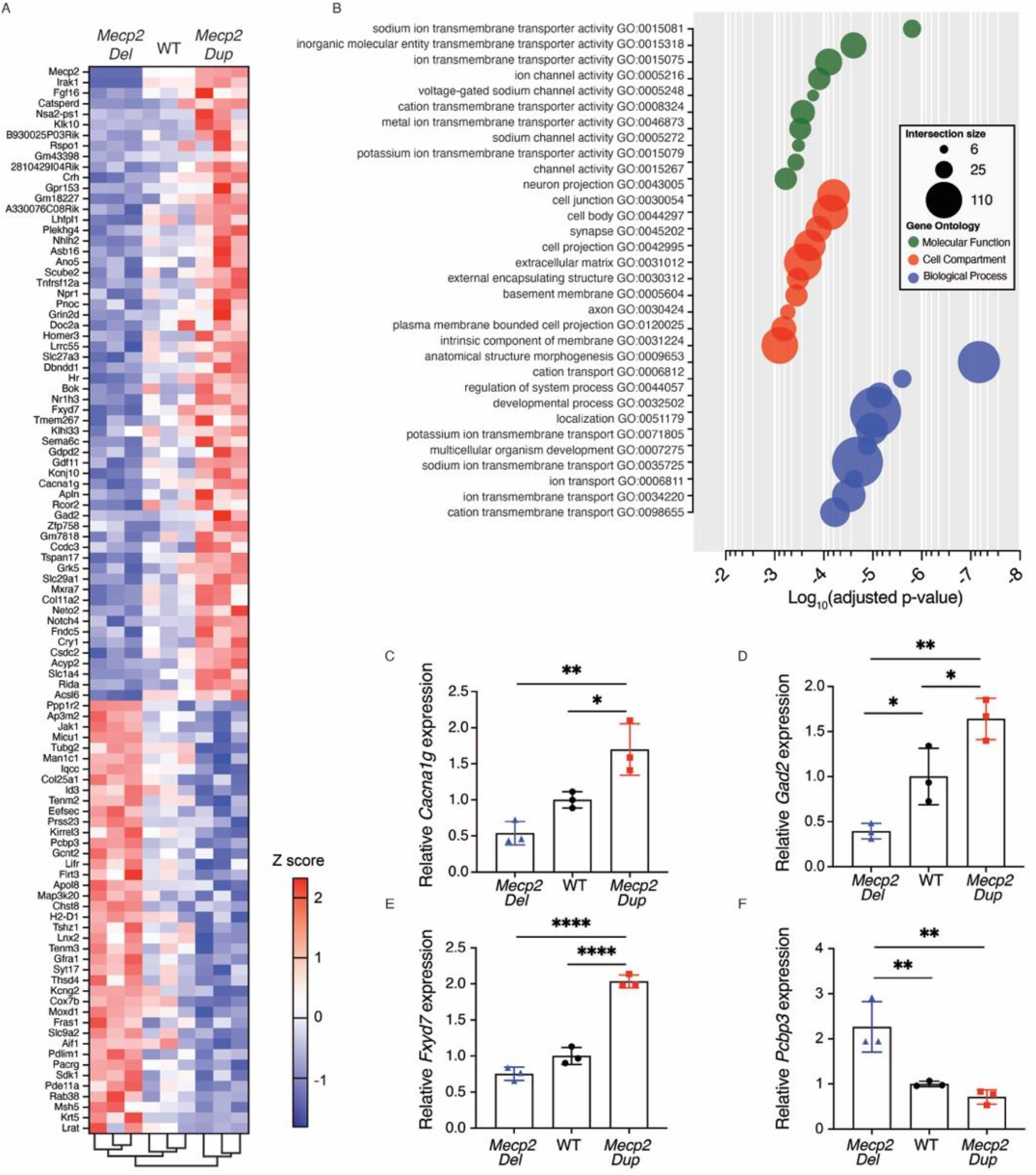
Altered gene expression in the hippocampus of *Mecp2 Dup* mice. **A)** Heat map of the top 100 differentially expressed genes in 6 weeks old *Mecp2 Dup* compared to *Mecp2 Del* mice and wild type controls. *Mecp2 Dup*, n=3; WT, n=3; *Mecp2 Del*, n=3. **B)** Gene ontology analysis identifying the 10 most enriched gene sets for molecular function, cell compartment and biological process. Analysis performed with g:Profiler (FDR<0.15%, base mean >10). RT-qPCR validated the expression levels of **C)** *Cacna1g*, **D)** *Gad2*, **E)** *Fxyd7*, **F)** *Pcpbp3* in *Mecp2 Dup* (n=3), WT (n=3), and *Mecp2 Del* mice (n=3). Statistical analysis was performed by one-way ANOVA followed by Tukey’s post hoc test. All data are represented as the mean +/− SD.

### *Mecp2 Dup* mice demonstrate an abnormal immune response with elevation in pro-inflammatory and Th1-associated cytokines and chemokines

Recurrent severe respiratory tract infections are a leading cause of morbidity and mortality among MDS patients, though the mechanisms underlying this phenomenon are incompletely understood. We infected mice with the *Influenza* H1N1 strain and assessed for immune abnormalities which would explain the heightened sensitivity of MDS patients to such infections. No difference in mortality monitored up to 14 days after infection was observed in *Mecp2 Dup* mice compared to wild type infected with the same dose of virus (Fig S8A). The percentage of body weight loss was similar between *Mecp2 Dup* and wild type mice (Fig S8B). Complete blood count and differential, and measurement of a panel of serum cytokines and chemokines were performed on days −1 (d.p.i. −1) and +4 (d.p.i. +4) following infection. Evaluation of cytokines and chemokines from bronchoalveolar lavage fluid (BALF) was performed on day +4.

At baseline on day −1, no notable differences were found between *Mecp2 Dup* mice and wildtype littermates both looking at blood cell counts and serum chemokines and cytokines panel (Fig 6A, Table S4). However, on day +4, elevated serum CCL5 (1.4-fold, p<0.01) and CXCL9 (1.7-fold, p<0.01) were noted (Fig 6D-E, Table S5). While interferon gamma (IFNγ) levels were elevated 2.23-fold in *Mecp2 Dup* mice, this was not statistically significant (p>0.999) (Fig 6C, Table S5). Moreover, blood lymphocytes were found increased in Mecp2 Dup mice compared to wild type littermates (1.96-fold, p<0.05) (Fig 6B). More robust alterations were identified in BALF, including increased levels of TNFα (2.19-fold, p<0.05), IFNγ (6.26-fold, p<0.05), CXCL9 (3.19-fold, p<0.01), CXCL10 (1.92-fold, p<0.05), the lymphocyte-recruiting chemokine CCL19 (1.60-fold, p<0.01), and IL-2 which promotes T-cell proliferation (1.69-fold, p<0.01) (Fig 6F, Table S6).

**Figure 6.**
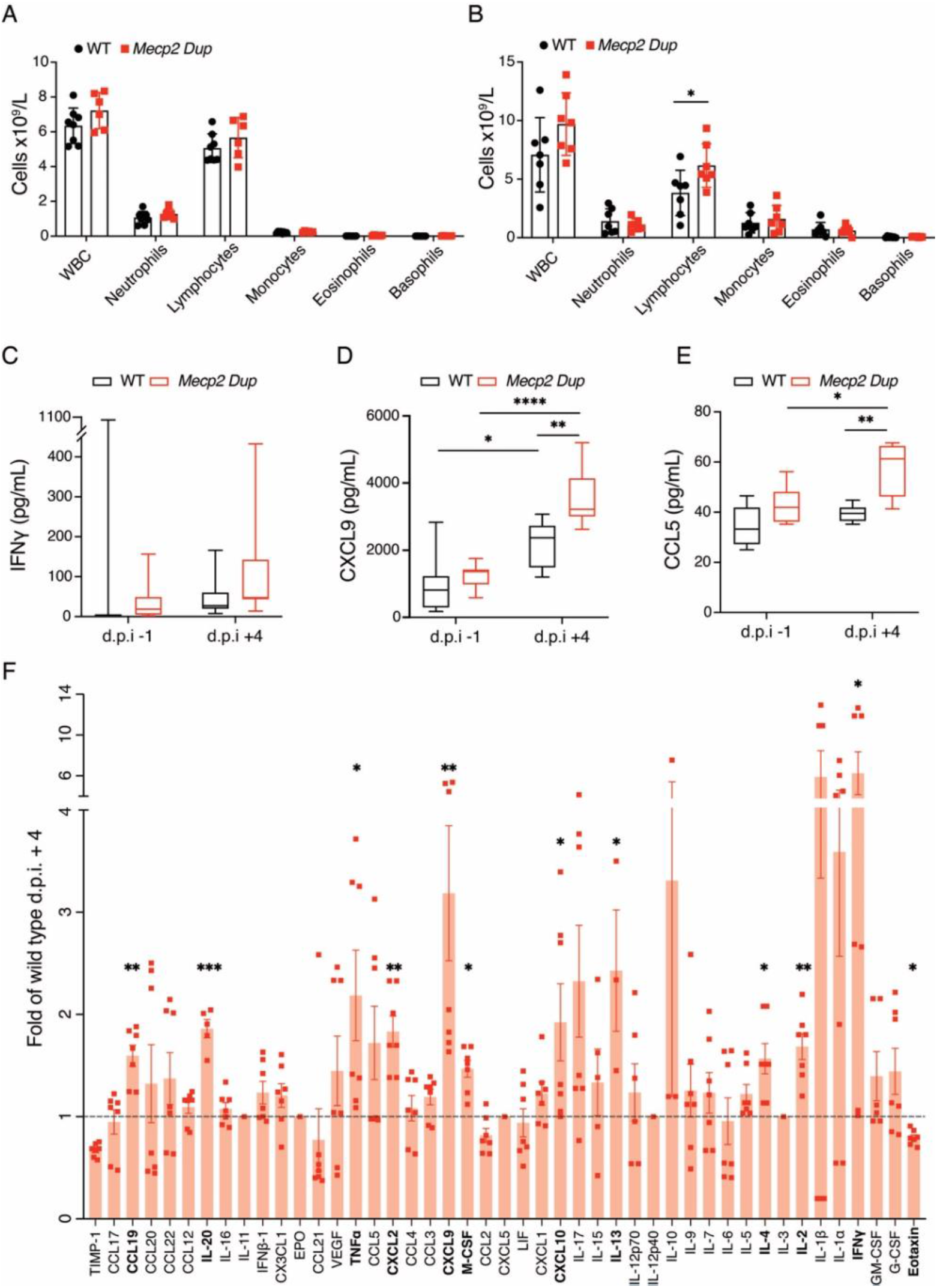
Abnormal immune response after influenza virus infection in the *Mecp2 Dup* mice. **A)** White blood cells count in naïve mice. *Mecp2 Dup*, n=6; WT, n=8. **B)** White blood cell count at day 4 after influenza infection. *Mecp2 Dup*, n=7; WT, n=7. Data are represented as the mean +/− SD. Statistical analyses were performed with Student’s t-test. Chemokines and cytokines multiplex analysis in the serum at day −1 before infection (d.p.i.-1) and day + 4 after influenza virus infection (d.p.i. +4) showed elevated production of **C)** IFNg, **D)** CCL5, and **E)** CXCL9. Data are represented as box plots, 5-95 percentile. Data was analyzed by one-way ANOVA followed by Tukey post hoc test or Kruskal Wallis test followed by Dunn’s multiple comparisons test. **F)** Cytokines and chemokines were measured in BALF. Data is presented as fold change versus wild type levels. *Mecp2 Dup*, n=7, WT, n=7. Data are represented as the mean +/− SEM. Statistical analyses were performed with Student’s t-test. * P<0.05, ** P<0.01, *** P<0.001, **** P<0.0001.

## Discussion

Modeling of complex genomic rearrangements remains a limiting factor in the study of disease-causing structural genomic variants in humans, of which MDS is a prime example. The current study showed the implementation of the fusion Cas9 proximity-based approach to successfully and efficiently generate an MDS mouse model harbouring a disease-defining 160kb tandem duplication involving both *Mecp2* and *Irak1* (*Mecp2 Dup*) and its reciprocal deletion mouse model (*Mecp2 Del*). *Mecp2 Dup* demonstrated abnormal neurodevelopmental features in keeping with patient phenotypes as well as existing MECP2-overexpression models. Moreover, *Mecp2 Dup* displayed some evidence for immune aberrations in keeping with enhanced inflammation and IFNγ activation, possibly owing to the contribution of heightened IRAK1 expression.

Efficient modelling of disease-causing CNVs is vital for the investigation of and therapeutic development in the context of rare inherited diseases. The current CRISPR/Cas9 dimerization approach increased the frequency of generating tandem duplications to 12% in mouse embryos, compared to the 1-3% efficiency previously observed by our group and others (Kraft et al., 2015; Maino et al., 2021; Pristyazhnyuk et al., 2019). This is likely due to the physical proximity of the rearranged DNA ends upon rapamycin treatment. This hypothesis is supported by previous work showing that binding of modified Cas9 to repair templates to ensure vicinity to the insertion site increased knock-in efficiencies (Gu et al., 2018; Ma et al., 2017; Savic et al., 2018). However, a more in-depth analysis would be needed to dissect the direct effect of the physical proximity on the duplication generation efficiency. Previous published studies investigated efficiencies of CNV generation by editing mESCs (Kraft et al., 2015; Maino et al., 2021), zygotes (Boroviak et al., 2016; Hara et al., 2016; Korablev et al., 2017) or one-cell stage embryos (Li et al., 2015). Instead, we performed the microinjection of the Cas9 mRNA and sgRNAs in 2-cell stage embryos that are known for having a longer G2 phase and open chromatin structure, and have previously been utilized to increase knock-in efficiencies (Gu et al., 2018). Moreover, we speculate that having two sister chromatids available might be critical in the case of duplication generation, where the second chromatid acts as a template for the generation of the tandem duplication. In addition, the inclusion of the bridging donor should favour the occurrence of minimal rearrangements at the duplication junction site.

The newly-generated *Mecp2 Dup* mouse model presents several possible advantages compared with transgenic models hitherto described. These include the faithful recapitulation of the genomic rearrangement seen in MDS patients, expression of *Mecp2* under its endogenous promoter, and the concurrent duplication and overexpression of *Irak1*. As anticipated, the *Mecp2 Dup* mouse model demonstrates neurodevelopmental phenotypes in keeping with previous observations from patients and transgenic mice. Prominent features include the involvement of hippocampi-related phenotypes in disease manifestations. As previously observed in the *MECP2-Tg* and *hDup* mice (Collins et al., 2004; Shao et al., 2021; Sztainberg et al., 2015), our MDS mouse model displays enhanced LTPs and increased contextual freezing behaviour, indicating abnormal synaptic plasticity and learning. In keeping with the latter, RNA-seq analysis was suggestive of dysregulated pathways of transmembrane ion transport which may lead to impaired neuronal excitability in the hippocampus. Differentially expressed genes included some well-known targets of MeCP2, such as *Gad2* and *Fxyd7*, together with genes recently found associated with intellectual disabilities and neurodevelopmental disorders, such as *Cacna1g* and *Pcpb3*. Another important disease manifestation was ataxia and motor coordination impairment in late disease stage, which has been reported in MDS patients (Ta et al., 2022) as well as the *Tau-Mecp2* mouse model (Na et al., 2012), and an increased brain-to-body ratio commonly noted in ASD patients and mouse models (Courchesne et al., 2019; Deliu et al., 2018). One feature noted in our mouse model but not in previous models is hyperactivity and reduced anxiety beginning at early disease stages. Such behaviours have been previously linked to intellectual disability in other mouse models, such as fragile X syndrome (Kazdoba et al., 2014). It is unclear why this feature characterizes our *Mecp2 Dup* mouse model, but not other MECP2-overexpression models.

Our *Mecp2 Dup* mouse model is the first to account for the minimal duplicated region shared across MDS patients, encompassing both the *Mecp2* and *Irak1* genes. Given the complex interplay reported between *Mecp2* and *Irak1*(Kishi et al., 2016), as well as their observed overexpression at the mRNA and protein level in the *Mecp2 Dup* mouse, it is possible the combined effects of both of these duplicated genes produce a different phenotype than that of MeCP2 overexpression alone. The variability in MDS disease phenotypes observed across various mouse lines could also be attributed to the different genetic backgrounds of the mice. In our case, we are modeling MDS on a CD-1 outbred background, while the other models were generated on inbred backgrounds such as C57BL/6 (Na et al., 2012), FVB (Collins et al., 2004), C57BL/6JOla × CBA hybrid (Koerner et al., 2018), and FVB/N × C57Bl/6 hybrid (Sztainberg et al., 2015).

Infections are a leading cause of morbidity and mortality in MDS patients. Recurrent infections have been reported in up to 75% of cases, with a predominance of respiratory disease caused by both viral and bacterial pathogens (Bauer et al., 2015). In this work, *Mecp2 Dup* mice showed no clear baseline differences in regard to major white blood cell subsets or serum cytokine and chemokine levels. Following *Influenza* infection, *Mecp2 Dup* mice developed elevated peripheral blood lymphocyte counts and aberrations in serum and BALF cytokines and chemokines suggestive of heightened T-cell proliferation (IL-2) and tissue recruitment (CCL19), as well as Th1 skewing (IFNγ, CXCL9, CXCL10, CCL5) and inflammation (TNT <2). The role of MECP2 in lymphocyte proliferation was previously demonstrated in cultured clonal T-cells from Rett syndrome patients, where MECP2 deficiency led to a growth disadvantage and a blunted response to mitogen stimulus (Balmer et al., 2002). Work in MDS patients previously identified an increase in naïve (CD45RA+) CD4+ T-cells, possibly also indicating an abnormal T-cell proliferation. Importantly, memory (CD45RO+) CD4+ T-cells were conversely reduced, suggestive of abnormal generation or reduced survival of memory T-cells (Bauer et al., 2015; Yang et al., 2012).

Our finding of Th1-skewing of the immune response following infection shows differences compared to findings in *MECP2*-overexpressing transgenic mice (Cronk et al., 2017; Yang et al., 2012). Yang et al reported that *Mecp2^Tg3^* mice, overexpressing up to 5 fold MeCP2 compared to wildtype, could not control an infection with the intra-macrophagic pathogen *Leishmania*, and showed impaired IFNγ secretion from involved lymph nodes. Their work also demonstrated decreased ability of CD4+ T-cells to differentiate into IFNγ-secreting cells *in vitro* (Yang et al., 2012). Subsequent work by Cronk et al., showed increased mortality of *Mecp2^Tg3^* mice following *Influenza A* infection, with decreased lung lymphocytes and reduced IFNγ in BALF (Cronk et al., 2017). However, from a clinical standpoint, MDS patients present quite differently from patients with complete or partial IFNγ-pathway deficiency, as the latter typically experience predisposition to infections with Mycobacteria*, Salmonella* sp. and intra-macrophagic pathogens (Bustamante et al., 2014), not seen in MDS patients (Bauer et al., 2015). One factor likely contributing to the discrepancy between our findings and those identified in *Mecp2^Tg3^* mice may be the role of IRAK1, overexpressed at both transcript and protein level in the *Mecp2 Dup* mice, which is unaccounted for by previous models. IRAK1 signaling was previously shown to cause Th1 differentiation via induction of T-bet (Cai et al., 2021; Deng et al., 2003; Heiseke et al., 2015). Additionally, increased IRAK1 expression in the context of Behcet’s disease was associated with increased production of IFNγ (Sun et al., 2018). Interestingly, even MeCP2 itself was previously demonstrated to be essential to differentiation of naïve CD4+ into Th1 cells (Jiang et al., 2014), while loss of MeCP2 expression in Rett syndrome patients was associated with a shift away from Th1 and loss of IFNγ expression. This suggests that perhaps the precise effect of MeCP2 aberrations could be dosedependent, whereby an overexpression up to 5-fold in leads to a different effect than a 2-fold increase. Clinically, chronic immune skewing toward Th1 has the potential to result in viral lung infections, chronic lung inflammation and impairment in immune memory, as noted for example in some STAT1 gain-of-function patients (Scott et al., 2022). However, further testing and sampling of both serum and tissue samples from acutely infected MDS patients as well as the Mecp2 Dup mice will be required to further address this issue.

From a treatment standpoint, The *Mecp2 Dup* mouse model could be instrumental to the testing and development of therapeutic strategies targeting MDS. Current MDS therapies in pre-clinical development are mainly based on the reduction of MECP2 expression levels by utilizing antisense oligonucleotides (ASOs) (Shao et al., 2021). The *Mecp2 Dup* model could be a valuable model for precise titration of ASOs to avoid phenotypic conversion to RTT. In addition, further ASOs could be designed and tested to target and ameliorate overexpression of other duplicated genes such as IRAK1. Moreover, specific gene editing strategies such as CRISPR/Cas9-based duplication correction (Maino et al., 2021) could be tested in the *Mecp2 Dup* model to precisely correct the disease-causing mutation and avoid the risk of over-targeting the duplicated genes.

In summary, our work showcases a platform to efficiently generate mouse models that faithfully recapitulate disease-causing structural variants. The newly-generated *Mecp2 Dup* mouse model provides an opportunity to model the neurodevelopmental traits of MDS patients, and further explore patient predisposition to severe respiratory disease via dysregulated interferon immunity. *Mecp2 Dup* mouse is a valuable tool to advance investigations of MDS disease mechanisms, as well as test and advance new genomic therapies for MDS patients.

## Author contributions

Conceptualization: E.M., B.G., E.A.I.; Methodology: E.M., O.S., S.Z.R, S.V., H.L., Y.Z., A.S.S; Investigation: E.M., O.S., S.Z.R, S.V., H.L., Y.Z.; Resources: R.D.C, B.G., J.R., E.A.I; Data curation: E.M., O.S., B.G., E.A.I.; Writing - original draft: E.M.; Writing - reviewing, and editing: E.M., O.S., S.Z.R., B.G., E.A.I., JR.; Supervision was by: M.W.S., Z.J., R.D.C, J.R., B.G., E.A.I.; Funding acquisition: E.M., R.D.C, J.R., B.G., E.A.I. All authors reviewed the final version of the manuscript.

## Conflict of interest

The authors declare no conflict of interest.

## Acknowledgments

We thank the members of the Cohn lab and Ivakine lab for their input in this study. We thank Marina Gertsenstein and Monica Pereira (Model Production Core, TCP) for their support with mouse model generation. We thank Sergio Pereira, Bhooma Thiruv, Tara Paton and Guillermo Casallo (The Centre for Applied Genomics) for their support with whole genome sequencing, RNA sequencing analysis, and ddPCR. We also thank Igor Vukobradovic and Zorana Berberovic (Clinical Phenotyping Core, TCP) for their assistance with mouse behavioural testing and blood cell count analysis. We thank Dr. Mitra Yousefi (Infection and Inflammation Core, TCP) for her support with influenza infection. This work was funded by the Reverse Rett Research Fund and Canadian Institutes of Health Research (CIHR (FDN-143334) to J.R.). Salary support for O.S. was provided by Ontario Ministry of Health Clinician Investigator Program, Canadian Child Health Clinician Scientist Program, Canadian Institutes of Health Research Canada Graduate Scholarship Doctoral Award, and the Hospital for Sick Children Clinician Scientist Training Program.

## Notes

### Competing Interest Statement

The authors have declared no competing interest.

