## Supplementary figures and tables for "A Cas9-fusion proximity-based approach generates an *Irak1-Mecp2* tandem duplication mouse model for the study of MeCP2 duplication syndrome"

**Figure S1.** Validation of the WGS results via Sanger sequencing.

**Figure S2.** The *Mecp2 Del* mouse model recapitulates RTT disease phenotypes.

**Figure S3.** *Mecp2* and *Irak1* are overexpressed in the cortex and cerebellum of *Mecp2 Dup* mice.

**Figure S4.** The *Mecp2 Dup* mice show increased activity and decreased anxiety in the centre of the open field arena.

**Figure S5.** The *Mecp2 Dup* mice show minimal motor coordination deficits in early disease stage.

**Figure S6.** Body weight analysis shows that *Mecp2 Dup* mice lose weight in late disease stages.

**Figure S7.** The *Mecp2 Dup* mice display a trend toward increased freezing behaviour on cued test.

**Figure S8.** Body weight and survival after influenza infection show no difference between *Mecp2 Dup* and wild type mice.

#### Supplementary figures

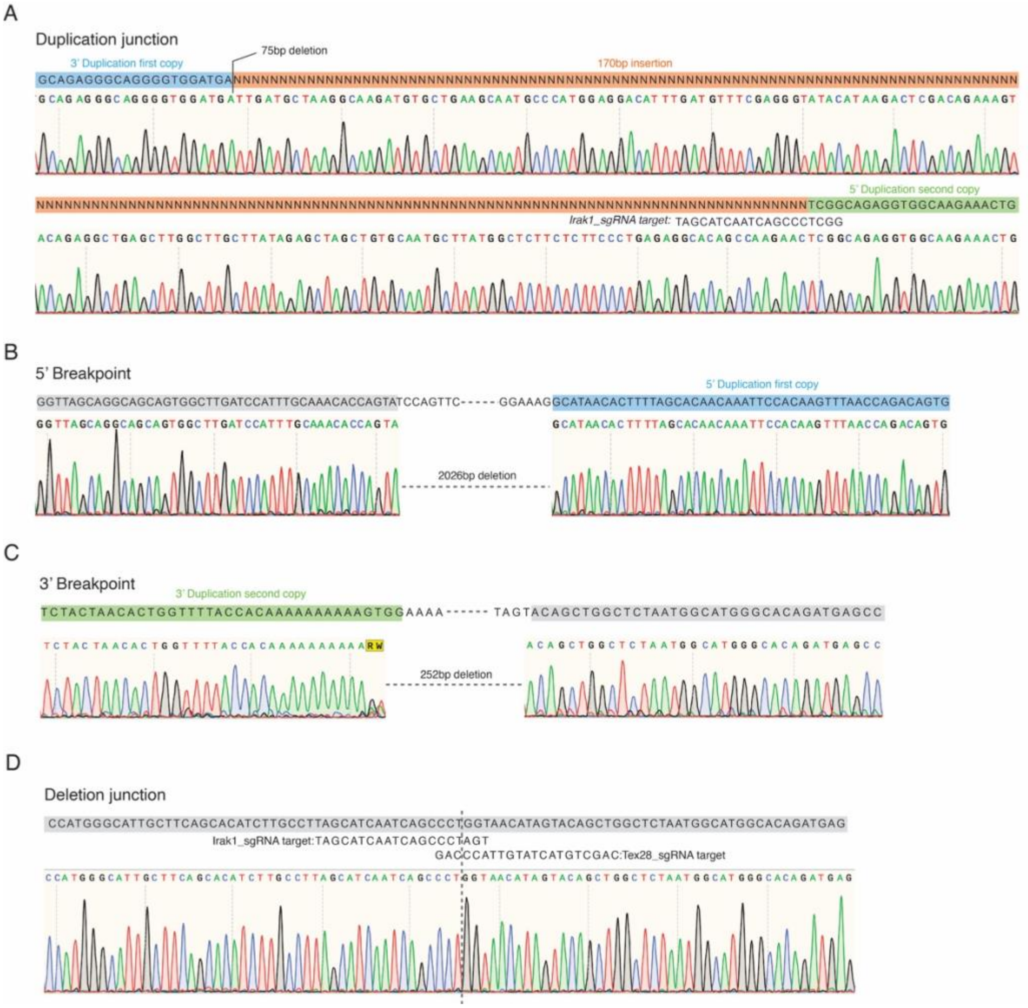


**Figure S1. Validation of the WGS results via Sanger sequencing. A)** Sanger sequencing of the duplication junction confirms the insertion of 170 bp at the junction site in the *Mecp2 Dup* mouse model. Breakpoints' sequencing shows presence of **B)** a 2026 bp deletion at the 5' breakpoint, around the Irak1_sgRNA target site, and **C)** a 252 bp deletion at the 3' breakpoint, around the Tex28_sgRNA target site. **D)** The *Mecp2 Del* mouse model presents a precise deletion junction.

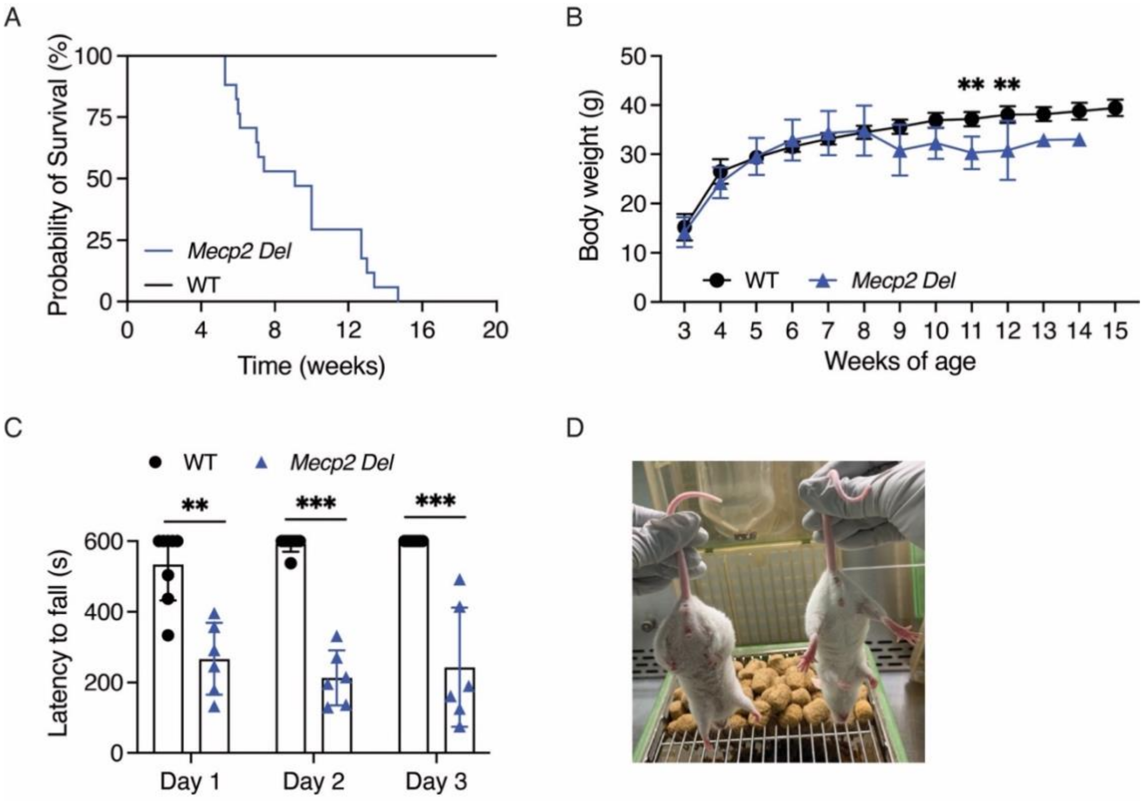


**Figure S2. The *Mecp2 Del* mouse model recapitulates RTT disease phenotypes. A)** *Mecp2 Del* mice showed reduced survival compared to wild type littermates. *Mecp2 Dup*, n=17; WT, n=11. Survival curves were compared with the Mantel-Cox test. *Mecp Del* median survival age = 9.1 weeks; P <0.0001. **B)** Body weight curve of the mice shown in A. Statistical analysis was performed by mixed-effect two-way ANOVA repeated measures followed by Bonferroni's multiple comparison test. **C)** Three-day rotarod test showed reduced motor coordination in the *Mecp2 Del* mice. *Mecp2 Del*, n=6; WT, n=8. Data was analyzed by using the using two-tailed Mann–Whitney *U* test. ** P <0.1, *** P<0.001. **D)** *Mecp2 Del* mice (Left) show typical RTT clasping behaviour. A wild type littermate (right) is utilized for comparison. Data are represented as the mean +/- SD.

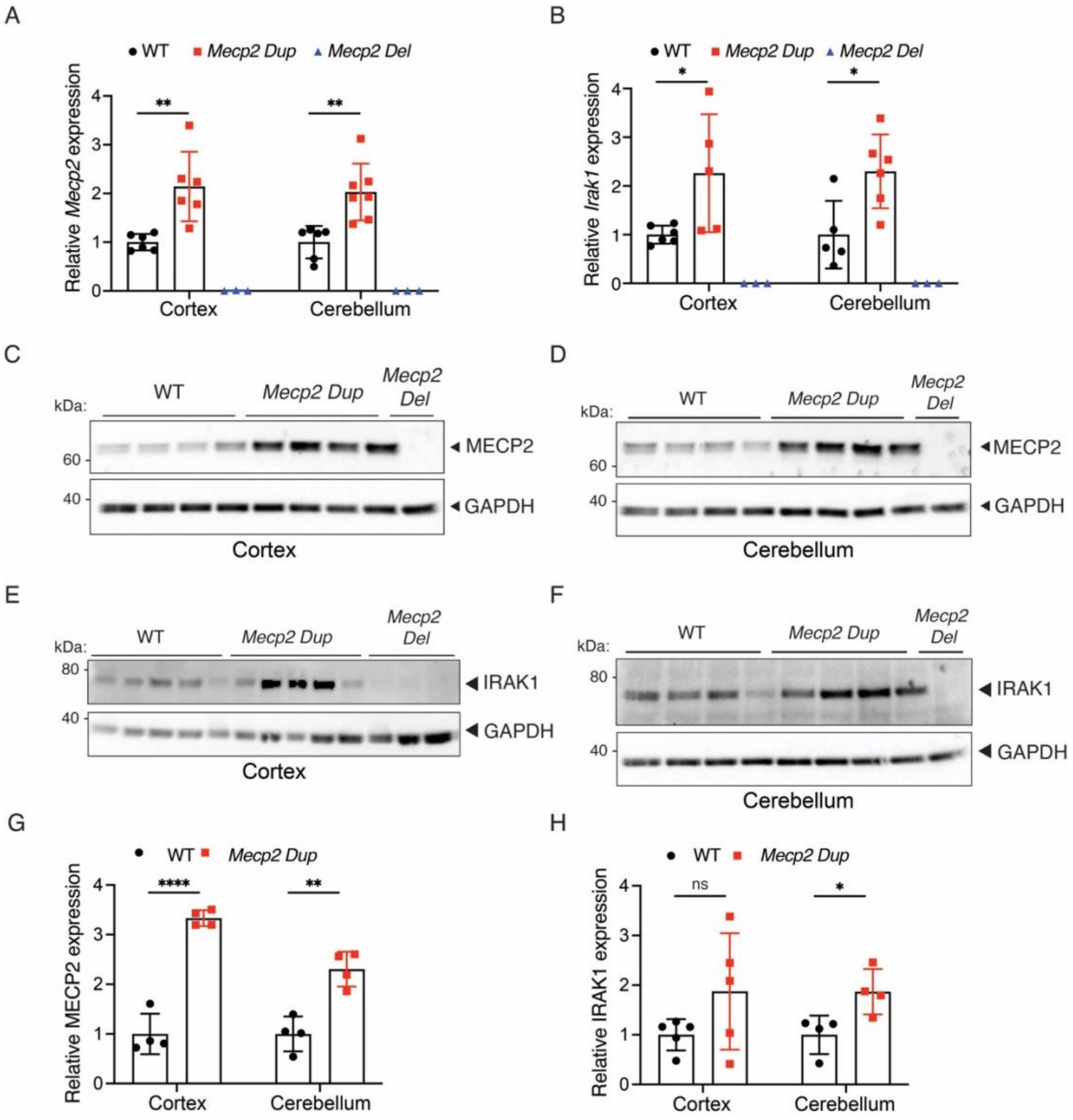


**Figure S3. *Mecp2* and *Irak1* are overexpressed in the cortex and cerebellum of *Mecp2 Dup* mice. A)** The levels of *Mecp2* expression were analyzed via qPCR in the cortex and cerebellum of 10 weeks *Mecp2 Dup* mice, wild type littermates and *Mecp2 Del* mice. The data is normalized over *Gapdh* expression. WT, n=5-6; *Mecp2 Dup*, n=6-7; *Mecp2 Del*, n=3. **B)** *Irak1* transcript level was analyzed via qPCR. The data is normalized over *Gapdh* expression. WT, n=5-6; *Mecp2 Dup*, n=5-6; *Mecp2 Del*, n=3. Western blot analysis confirmed increase of MECP2 expression **C)** in the cortex and **D)** in the cerebellum of *Mecp2 Dup* mice compared to wild type littermates. GAPDH serves as a loading control. IRAK1 expression was analyzed via Western Blot in **E)** cortex and **F)** cerebellum. GAPDH serves as a loading control. Densitometry analysis to quantify the amount of **G)** MECP2 and **H)** IRAK1 expression in various brain areas. WT, n=4-5; *Mecp2 Dup*, n=4-5; *Mecp2 Del*, n=1-3. All data are represented as the mean +/- SD. Statistical analyses were performed with Student's t-test. *P<0.5, **P<0.01, ****P<0.0001.

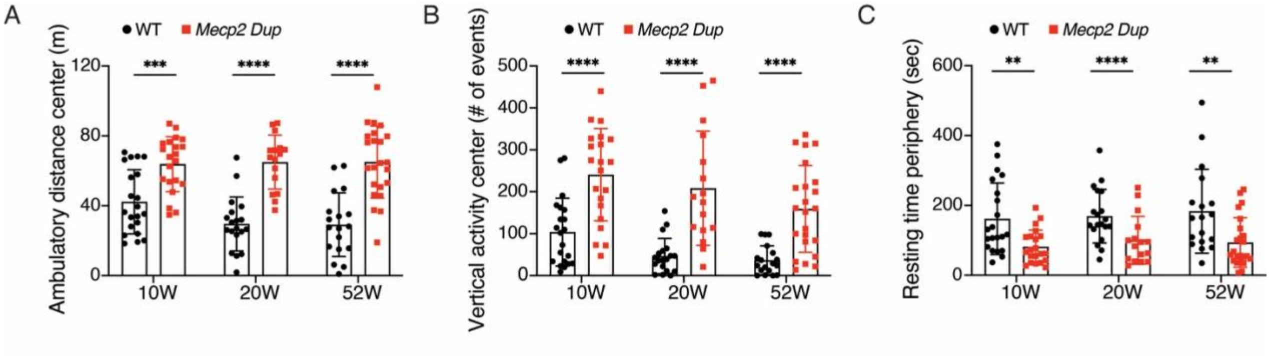


**Figure S4. The *Mecp2 Dup* mice show increased activity and decreased anxiety in the centre of the open field arena.** The activity of *Mecp2 Dup* mice compared to wild type littermates was evaluated at 10, 20 and 52 weeks in the open field arena. All parameters analyzed in centre of the arena were highly dysregulated compared to wild type mice. **A)** Ambulatory distance in the center, **B)** Vertical activity in the center and **C)** Resting time in the periphery were measured. *Mecp2 Dup*, n=16-23; WT, n=18-21. Statistical analyses were performed with Student's t-test for data normally distributed or Mann–Whitney *U* test for data that do not follow a normal distribution. All data are represented as the mean +/- SD. **P<0.01, ***P<0.001, **** P<0.0001.

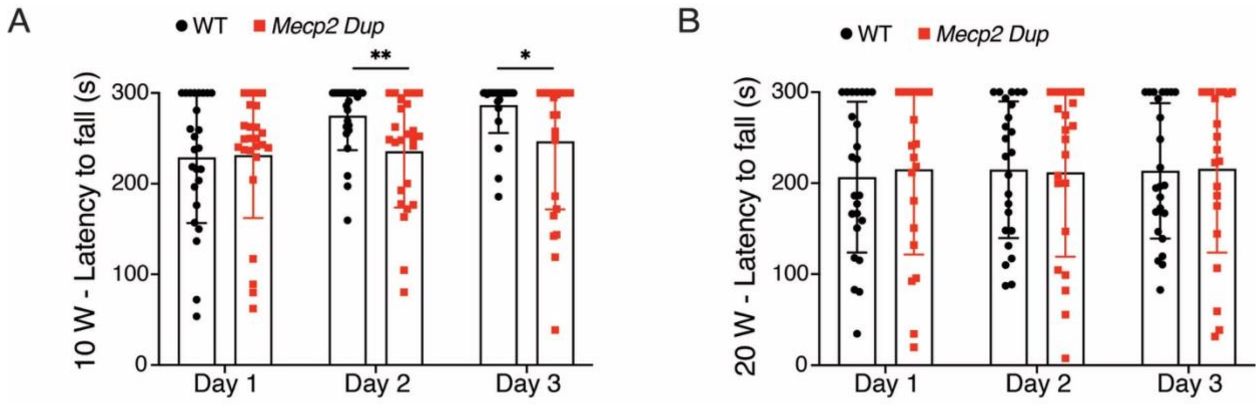


**Figure S5. The *Mecp2 Dup* mice show minimal motor coordination deficits in early disease stage.** A 3 days rotarod test performed to evaluate motor coordination of the *Mecp2 Dup mice* at **A)** 10 weeks of age and **B)** 20 weeks of age. Small differences were observed among the two groups. *Mecp2 Dup*, n=24-25; WT, n=21-25. Statistical analyses were performed with the Mann–Whitney *U* test or Student's t-test for data normally distributed. All data are represented as the mean +/- SD. *P<0.5, **P<0.01.

A

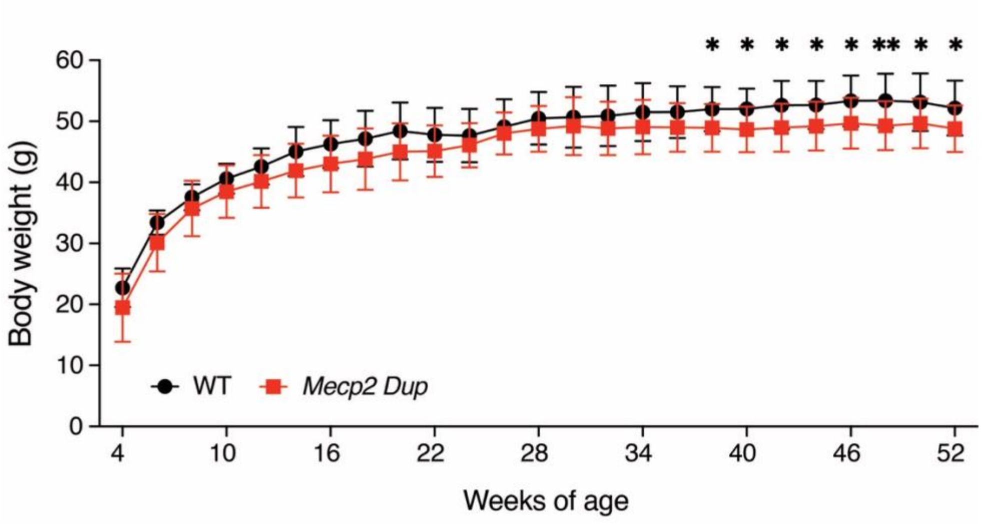


**Figure S6. Body weight analysis shows that *Mecp2 Dup* mice lose weight in late disease stages. A)** Body weight was monitored weekly up from 4 to 52 weeks of age. *Mecp2 Dup*, n=11-17; WT, n=11-12. Statistical analysis was performed by mixed-effect two-way ANOVA repeated measures followed by Bonferroni's multiple comparison test. All data are represented as the mean +/- SD.

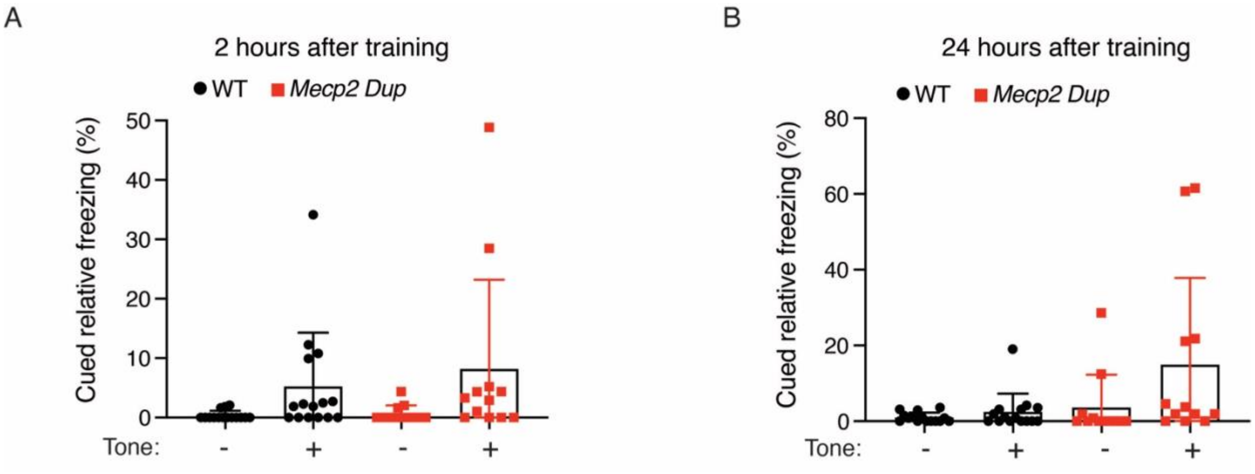


**Figure S7. The *Mecp2 Dup* mice display a trend toward increased freezing behaviour on cued test. A)** No significant changes in percent freezing were shown when cued freezing behaviour was assessed both at 2 hours and **B)** 24 hours after training. *Mecp2 Dup,* n=12; WT, n=15. Statistical analysis performed by Kruskal Wallis test followed by Dunn's multiple comparison test. All data are represented as the mean +/- SD.

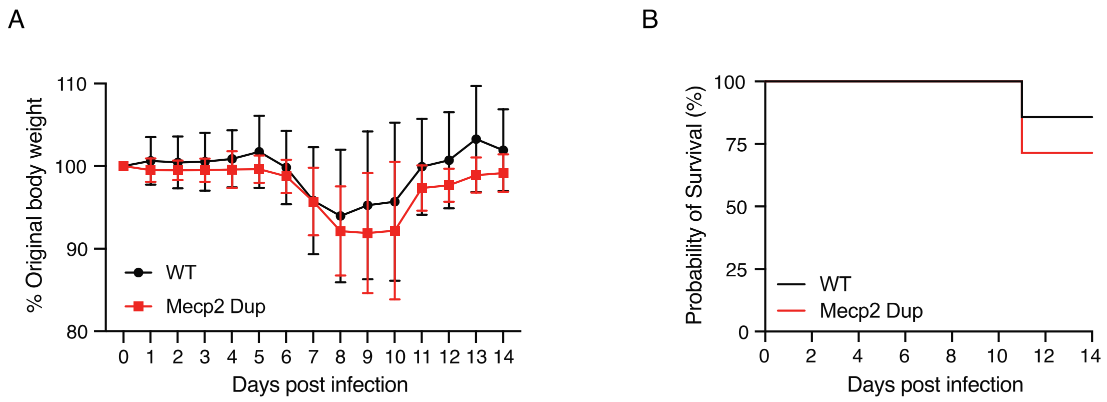


**Figure S8. Body weight and survival after influenza infection show no difference between *Mecp2 Dup* and wild type mice.** **A)** Weight loss upon infection in *Mecp2 Dup* and wild type littermates was measured up to day 14 post infection as percentage of body weight compared to day 0. *Mecp2 Dup*, n=5-7; wild type, n=6-7. Statistical analysis was performed by mixed-effect two-way ANOVA repeated measures followed by Bonferroni's multiple comparison test. Data are represented as the mean +/- SD. **B)** Survival of 10 weeks old *Mecp2 Dup* and wild type littermates infected with a dose of influenza shows no differences up to day 14 post infection. *Mecp2 Dup*, n=7; wild type, n=7. Statistical analysis performed with Mantel-Cox test (P = 0.5302).

**Supplementary tables**

**Table S1.** *Mecp2 Dup* – WGS structural variants.

**Table S2.** RNA-seq differentially expressed genes between *Mecp2 Dup* and *Mecp2 Del* mice. (Provided as excel)

**Table S3.** Gene ontology analysis. (Provided as excel)

**Table S4.** Cytokine/Chemokine 44-Plex Discovery Assay® Array of Serum of *Mecp2 Dup* and wild type mice the day prior (d.p.i -1) to *Influenza* infection.

**Table S5.** Cytokine/Chemokine 44-Plex Discovery Assay® Array of Serum of *Mecp2 Dup* and wild type mice at day 4 post *Influenza* infection (d.p.i +4).

**Table S6.** Cytokine/Chemokine 44-Plex Discovery Assay® Array of BALF of *Mecp2 Dup* and wild type mice at day 4 post *Influenza* infection (d.p.i +4).

**Table S7.** sgRNAs and bridging donor utilized to generate the *Mecp2 Dup* mice.

**Table S8.** Oligonucleotides utilized in this study.

**Table S1.** ***Mecp2 Dup*** **– WGS structural variants**

| **Chr** | **Pos start** | **Pos end** | **SV length (bp)** | **SV type** | **What** |
|---|---|---|---|---|---|
| X | 74,012,223 | 74,014,249 | -2026 | DEL | 5’ breakpoint deletion |
| X | 74,014,249 | 74,172,116 | 157,876 | DUP | *Irak1-Tex28* dup |
| X | 74,012,505 | 74,012,335 | 170 | INS | Insertion at the duplication junction |
| X | 74,171,949 | 74,172,201 | -252 | DEL | 3’ breakpoint deletion |

**Table S2. RNA-seq differentially expressed genes between *Mecp2 Dup* and *Mecp2 Del* mice**

| Gene Name | Adjusted p value | log2 Fold Change |
|---|---|---|
| Mecp2 | 2.11E-145 | 9.70 |
| Irak1 | 3.29E-104 | 8.92 |
| Pgr15l | 0.1309 | 6.13 |
| A830036E02Rik | 0.0494 | 5.49 |
| Lhx5 | 0.0719 | 4.67 |
| BC035947 | 0.0726 | 4.30 |
| Fgf16 | 0.0005 | 4.18 |
| AW551984 | 0.0753 | 4.09 |
| H2-Q2 | 0.0874 | 3.71 |
| Chodl | 0.1119 | 3.65 |
| Lemd1 | 0.0524 | 3.29 |
| Catsperd | 0.0102 | 3.29 |
| Smyd1 | 0.0748 | 3.26 |
| Nsa2-ps1 | 0.0309 | 3.07 |
| Klk10 | 4.05E-06 | 3.00 |
| B930025P03Rik | 0.0006 | 2.92 |
| 1110015O18Rik | 0.1225 | 2.73 |
| Nxph4 | 0.1000 | 2.67 |
| Abhd11os | 0.0719 | 2.61 |
| Prph | 0.0970 | 2.58 |
| Oprk1 | 0.1260 | 2.55 |
| Rspo1 | 0.0237 | 2.42 |
| Gm43398 | 0.0066 | 2.41 |
| Hmcn2 | 0.1110 | 2.40 |
| 2810429I04Rik | 0.0066 | 2.35 |
| Slc5a7 | 0.1390 | 2.33 |
| Gm29695 | 0.0701 | 2.33 |
| Crh | 0.0005 | 2.32 |
| Gm21860 | 0.1315 | 2.32 |
| Frem3 | 0.1246 | 2.31 |
| 9230009I02Rik | 0.0524 | 2.12 |
| Slc18a2 | 0.0327 | 2.08 |
| Prkcd | 0.0371 | 2.08 |
| Cngb1 | 0.1176 | 2.08 |
| Gpr153 | 2.35E-05 | 2.03 |

| | | |
|---|---|---|
| Spn | 0.1286 | 2.00 |
| Scn9a | 0.0847 | 1.98 |
| Gm18227 | 0.0187 | 1.97 |
| H1f3 | 0.0493 | 1.95 |
| A330076C08Rik | 0.0262 | 1.93 |
| Rnase1 | 0.0672 | 1.92 |
| Lhfpl1 | 0.0100 | 1.87 |
| BC065397 | 0.0434 | 1.87 |
| Rab37 | 0.0922 | 1.80 |
| Grid2ip | 0.0673 | 1.70 |
| Plekhg4 | 0.0017 | 1.68 |
| Nhlh2 | 0.0002 | 1.68 |
| 4930412C18Rik | 0.0922 | 1.65 |
| Cpne9 | 0.0533 | 1.57 |
| Asb16 | 0.0038 | 1.56 |
| Mctp2 | 0.1133 | 1.56 |
| Ret | 0.0769 | 1.55 |
| Cyp2s1 | 0.0434 | 1.49 |
| Zdhhc22 | 0.0691 | 1.48 |
| Cdh23 | 0.0434 | 1.47 |
| Hcn4 | 0.1460 | 1.47 |
| Ano5 | 0.0174 | 1.42 |
| Tchh | 0.0347 | 1.40 |
| Gbp4 | 0.0451 | 1.38 |
| Sptssb | 0.1373 | 1.35 |
| Cyp26b1 | 0.1489 | 1.34 |
| Scube2 | 0.0285 | 1.32 |
| Tnfrsf12a | 0.0013 | 1.31 |
| Mafa | 0.1176 | 1.30 |
| Npr1 | 0.0133 | 1.28 |
| Gm10863 | 0.0391 | 1.28 |
| St8sia6 | 0.0582 | 1.28 |
| Vwc2 | 0.0402 | 1.27 |
| Inhba | 0.0779 | 1.27 |
| Egfl6 | 0.1116 | 1.25 |
| Arg2 | 0.0886 | 1.25 |
| Penk | 0.1451 | 1.23 |
| Mill2 | 0.0545 | 1.22 |

| | | |
|---|---|---|
| Pnoc | 0.0284 | 1.21 |
| Vat1 | 0.1456 | 1.21 |
| Grin2d | 0.0206 | 1.18 |
| Doc2a | 0.0309 | 1.17 |
| Homer3 | 0.0009 | 1.16 |
| Lrrc55 | 0.0009 | 1.13 |
| Iigp1 | 0.0559 | 1.13 |
| B430212C06Rik | 0.1221 | 1.12 |
| Adora2b | 0.1373 | 1.12 |
| Slc27a3 | 0.0006 | 1.11 |
| Dbndd1 | 0.0005 | 1.09 |
| Tmem196 | 0.0362 | 1.08 |
| Zfp605 | 0.1433 | 1.07 |
| Hr | 1.88E-06 | 1.07 |
| Stc1 | 0.0709 | 1.06 |
| Bok | 0.0077 | 1.06 |
| Nr1h3 | 0.0200 | 1.06 |
| Klk8 | 0.0942 | 1.06 |
| Fxyd7 | 4.05E-06 | 1.05 |
| Tmem267 | 0.0102 | 1.04 |
| Cdh24 | 0.1119 | 1.03 |
| Klhl33 | 0.0185 | 1.02 |
| Sema6c | 0.0001 | 1.00 |
| Aldh1a1 | 0.0964 | 1.00 |
| Gdpd2 | 0.0125 | 0.98 |
| Gdf11 | 6.09E-05 | 0.98 |
| Plcxd2 | 0.0434 | 0.97 |
| Gm37691 | 0.1337 | 0.96 |
| Rnd1 | 0.1246 | 0.94 |
| Kcnj10 | 2.90E-09 | 0.94 |
| Klhl13 | 0.0533 | 0.91 |
| Kcnk4 | 0.0886 | 0.90 |
| Cast | 0.0922 | 0.90 |
| Cacna1g | 2.62E-05 | 0.88 |
| Apln | 0.0017 | 0.87 |
| Rcor2 | 0.0133 | 0.86 |
| Gad2 | 0.0309 | 0.85 |
| Zfp758 | 0.0248 | 0.83 |

| | | |
|---|---|---|
| Gm7818 | 0.0187 | 0.82 |
| Ighm | 0.0332 | 0.81 |
| Lpl | 0.0494 | 0.80 |
| Asic4 | 0.1225 | 0.80 |
| Ccdc3 | 0.0232 | 0.77 |
| Serpinb8 | 0.1428 | 0.76 |
| Nrep | 0.0669 | 0.76 |
| Kcnip1 | 0.0645 | 0.76 |
| Tspan17 | 3.01E-09 | 0.75 |
| Abcb10 | 0.0708 | 0.74 |
| Trim67 | 0.0919 | 0.74 |
| Adamts6 | 0.0326 | 0.74 |
| Lrrc75b | 0.0591 | 0.73 |
| Cidea | 0.0769 | 0.72 |
| Hcn3 | 0.1460 | 0.71 |
| Pkib | 0.0768 | 0.71 |
| Slc9a5 | 0.1311 | 0.70 |
| Grk5 | 0.0285 | 0.70 |
| Gpr83 | 0.0581 | 0.69 |
| Rbpj | 0.0380 | 0.69 |
| Tmem158 | 0.0451 | 0.68 |
| Slc29a1 | 0.0017 | 0.68 |
| A2ml1 | 0.1225 | 0.67 |
| Nudt12 | 0.1221 | 0.66 |
| Sgsm1 | 0.0756 | 0.66 |
| Mxra7 | 0.0237 | 0.63 |
| Col11a2 | 0.0040 | 0.63 |
| Trhde | 0.0563 | 0.63 |
| Neto2 | 0.0006 | 0.62 |
| Notch4 | 0.0229 | 0.62 |
| Mir670hg | 0.0885 | 0.61 |
| Gm15328 | 0.1451 | 0.61 |
| Zfp40 | 0.0340 | 0.61 |
| Nrarp | 0.1337 | 0.61 |
| Trib2 | 0.0977 | 0.61 |
| Sass6 | 0.1260 | 0.60 |
| Samd10 | 0.1273 | 0.59 |
| Pcp4l1 | 0.1095 | 0.57 |

| | | |
|---|---|---|
| Fndc5 | 0.0048 | 0.57 |
| Cry1 | 0.0212 | 0.56 |
| Gm45495 | 0.1326 | 0.55 |
| Rflnb | 0.1119 | 0.54 |
| Rragd | 0.0615 | 0.53 |
| Csdc2 | 1.96E-05 | 0.53 |
| Insig1 | 0.0411 | 0.52 |
| Gnal | 0.0559 | 0.52 |
| Acyp2 | 0.0237 | 0.49 |
| Spock3 | 0.1360 | 0.48 |
| Slc1a4 | 0.0020 | 0.48 |
| Nes | 0.1441 | 0.47 |
| Rrp1b | 0.1463 | 0.47 |
| Prkx | 0.0607 | 0.47 |
| Tmem229a | 0.0756 | 0.46 |
| Zfp946 | 0.0754 | 0.45 |
| Itgb8 | 0.0752 | 0.45 |
| Adal | 0.1139 | 0.44 |
| Arxes1 | 0.0657 | 0.44 |
| Metrn | 0.1432 | 0.44 |
| Vwa1 | 0.0451 | 0.44 |
| Jam2 | 0.0906 | 0.42 |
| Atp1b2 | 0.1000 | 0.42 |
| Hmgn5 | 0.1041 | 0.42 |
| Cmpk2 | 0.0774 | 0.42 |
| Sat1 | 0.0769 | 0.39 |
| Wnk3 | 0.0939 | 0.39 |
| Zcchc24 | 0.1464 | 0.39 |
| Dner | 0.0340 | 0.38 |
| Rala | 0.0769 | 0.38 |
| Nefm | 0.1125 | 0.38 |
| Tmem62 | 0.1056 | 0.37 |
| Rida | 0.0243 | 0.37 |
| Gm12258 | 0.0922 | 0.35 |
| Tppp | 0.0465 | 0.35 |
| Tango2 | 0.1282 | 0.35 |
| Tagln3 | 0.1233 | 0.35 |
| Cygb | 0.1110 | 0.34 |

| | | |
|---|---|---|
| Chst10 | 0.0989 | 0.34 |
| Acsl6 | 0.0057 | 0.33 |
| Aldoc | 0.1176 | 0.32 |
| Nfic | 0.1311 | 0.31 |
| 2610301B20Rik | 0.0850 | 0.30 |
| Stmn2 | 0.1233 | 0.30 |
| Tmem263 | 0.0779 | 0.30 |
| Ddhd1 | 0.0756 | 0.29 |
| Hcn2 | 0.1167 | 0.29 |
| Zyx | 0.1326 | 0.29 |
| Gpr158 | 0.0756 | 0.29 |
| Ttyh1 | 0.1459 | 0.26 |
| Mapk8ip1 | 0.0836 | 0.25 |
| Dcaf7 | 0.0859 | -0.24 |
| Eno1b | 0.1347 | -0.25 |
| Rtn1 | 0.1313 | -0.26 |
| Ppp1r2 | 0.0187 | -0.30 |
| Ap3m2 | 0.0102 | -0.31 |
| Ube2e3 | 0.1119 | -0.33 |
| Ppp1r9a | 0.1159 | -0.34 |
| Ddx39b | 0.0347 | -0.34 |
| Cep83os | 0.0649 | -0.35 |
| Tmem94 | 0.1246 | -0.35 |
| Atp6v1g2 | 0.0510 | -0.36 |
| Nfia | 0.0410 | -0.36 |
| Zfp316 | 0.1246 | -0.37 |
| Rps6ka2 | 0.0559 | -0.39 |
| Ctdspl | 0.1010 | -0.39 |
| Rpa1 | 0.0981 | -0.39 |
| Dennd1b | 0.0456 | -0.39 |
| Jak1 | 0.0059 | -0.40 |
| Triqk | 0.1393 | -0.40 |
| Ripor2 | 0.0388 | -0.41 |
| Kat14 | 0.0473 | -0.41 |
| Brpf3 | 0.0559 | -0.41 |
| Lrrc8d | 0.0808 | -0.42 |
| Eef2k | 0.0451 | -0.42 |
| Coq8a | 0.0989 | -0.42 |

| | | |
|---|---|---|
| Csgalnact1 | 0.0841 | -0.43 |
| Micu1 | 0.0019 | -0.45 |
| Tubg2 | 0.0040 | -0.45 |
| Fjx1 | 0.1402 | -0.46 |
| Trp53bp2 | 0.1221 | -0.46 |
| Rgs12 | 0.0981 | -0.47 |
| Asic2 | 0.0361 | -0.47 |
| Nrip1 | 0.0964 | -0.47 |
| Atg101 | 0.0434 | -0.47 |
| Flna | 0.1419 | -0.50 |
| Top1mt | 0.0989 | -0.50 |
| Tle3 | 0.0989 | -0.50 |
| Col4a2 | 0.0327 | -0.52 |
| Man1c1 | 0.0179 | -0.52 |
| Sgk1 | 0.1119 | -0.53 |
| Arhgef28 | 0.0801 | -0.54 |
| Iqcc | 0.0029 | -0.54 |
| Naaa | 0.0434 | -0.54 |
| Firre | 0.0483 | -0.55 |
| Mterf2 | 0.0906 | -0.56 |
| Septin9 | 0.0665 | -0.57 |
| Col25a1 | 0.0065 | -0.58 |
| Id3 | 0.0297 | -0.58 |
| Col4a1 | 0.0340 | -0.59 |
| Foxo1 | 0.0770 | -0.59 |
| Bag3 | 0.0998 | -0.60 |
| Slc24a4 | 0.0897 | -0.60 |
| Tenm2 | 0.0008 | -0.61 |
| Mrpl33 | 0.1070 | -0.61 |
| St6galnac5 | 0.0310 | -0.61 |
| Plch2 | 0.0623 | -0.61 |
| Eefsec | 0.0125 | -0.62 |
| Igfbp4 | 0.0847 | -0.62 |
| Prss23 | 0.0034 | -0.63 |
| Ap1g2 | 0.0312 | -0.63 |
| Foxp1 | 0.1104 | -0.63 |
| Kirrel3 | 0.0016 | -0.64 |
| Pde1a | 0.0989 | -0.65 |

| | | |
|---|---|---|
| Pcbp3 | 2.16E-09 | -0.67 |
| Acvr1c | 0.0977 | -0.68 |
| Gm26871 | 0.1040 | -0.68 |
| Lmo4 | 0.0427 | -0.68 |
| Klf2 | 0.1402 | -0.68 |
| Gcnt2 | 0.0102 | -0.68 |
| Itga4 | 0.1471 | -0.69 |
| Fbn1 | 0.1452 | -0.69 |
| Smad3 | 0.0434 | -0.69 |
| Gm7694 | 0.0654 | -0.70 |
| Lifr | 0.0285 | -0.72 |
| Pygm | 0.0711 | -0.74 |
| Gsg1l | 0.1246 | -0.74 |
| Flrt3 | 0.0197 | -0.75 |
| Fam81a | 0.0791 | -0.75 |
| Myo16 | 0.0836 | -0.77 |
| Greb1l | 0.0752 | -0.77 |
| Apol8 | 0.0225 | -0.77 |
| Map3k20 | 0.0243 | -0.78 |
| Eya4 | 0.0847 | -0.78 |
| Chst8 | 1.88E-05 | -0.79 |
| H2-D1 | 0.0075 | -0.79 |
| Crhbp | 0.0628 | -0.80 |
| Gm26644 | 0.0340 | -0.82 |
| 5730522E02Rik | 0.1334 | -0.84 |
| Tshz1 | 0.0055 | -0.84 |
| Lnx2 | 0.0200 | -0.86 |
| Tenm3 | 0.0017 | -0.87 |
| Marchf3 | 0.0752 | -0.90 |
| Tspan9 | 0.0451 | -0.91 |
| 9030407P20Rik | 0.0900 | -0.91 |
| Gfra1 | 6.87E-09 | -0.92 |
| Upp2 | 0.1440 | -0.93 |
| Etv1 | 0.0836 | -0.94 |
| Krt73 | 0.0550 | -1.03 |
| Syt17 | 0.0027 | -1.04 |
| Thsd4 | 0.0065 | -1.05 |
| Kcng2 | 0.0006 | -1.06 |

| | | |
|---|---|---|
| Crispld2 | 0.0977 | -1.06 |
| Cox7b | 0.0004 | -1.07 |
| Arhgap10 | 0.0974 | -1.07 |
| Moxd1 | 0.0002 | -1.08 |
| Fras1 | 0.0183 | -1.08 |
| Slc9a2 | 0.0005 | -1.11 |
| Aif1 | 0.0055 | -1.13 |
| Pdlim1 | 0.0018 | -1.14 |
| Gm19744 | 0.0434 | -1.15 |
| Gm14410 | 0.1243 | -1.17 |
| Wnt5b | 0.0922 | -1.18 |
| Olfml2a | 0.1334 | -1.18 |
| Ano2 | 0.1347 | -1.18 |
| Pacrg | 1.06E-06 | -1.20 |
| Gask1a | 0.0434 | -1.20 |
| Gm43174 | 0.0559 | -1.40 |
| Sdk1 | 0.0026 | -1.43 |
| Bmp3 | 0.0494 | -1.46 |
| Gm37107 | 0.0347 | -1.53 |
| Slc5a5 | 0.0451 | -1.61 |
| Col6a2 | 0.0984 | -1.65 |
| Twist1 | 0.1382 | -1.73 |
| Gm47026 | 0.1256 | -1.76 |
| Creb3l3 | 0.0974 | -1.95 |
| Rps3a2 | 0.1176 | -1.99 |
| Serpina3h | 0.0494 | -1.99 |
| Pde11a | 1.88E-05 | -2.02 |
| Clec18a | 0.0451 | -2.13 |
| Serpina3i | 0.1132 | -2.18 |
| Rab38 | 0.0111 | -2.24 |
| Col6a1 | 0.1119 | -2.47 |
| Msh5 | 0.0052 | -2.77 |
| Slc25a41 | 0.0340 | -2.84 |
| Gm14419 | 0.0667 | -3.02 |
| Krt5 | 0.0040 | -3.69 |
| Lrat | 0.0059 | -4.25 |
| Gm9847 | 0.0380 | -4.80 |
| Gm10654 | 0.0605 | -7.47 |

### Table S3. Gene ontology analysis

| Source | Name | ID | Adjusted p value | Gene set size | Intersection size |
|---|---|---|---|---|---|
| GO:MF | sodium ion transmembrane transporter activity | GO:0015081 | 1.59E-06 | 137 | 14 |
| GO:MF | inorganic molecular entity transmembrane transporter activity | GO:0015318 | 2.49E-05 | 699 | 29 |
| GO:MF | ion transmembrane transporter activity | GO:0015075 | 7.97E-05 | 826 | 31 |
| GO:MF | ion channel activity | GO:0005216 | 0.000124112 | 429 | 21 |
| GO:MF | voltage-gated sodium channel activity | GO:0005248 | 0.000165583 | 23 | 6 |
| GO:MF | cation transmembrane transporter activity | GO:0008324 | 0.000268831 | 611 | 25 |
| GO:MF | metal ion transmembrane transporter activity | GO:0046873 | 0.000303618 | 415 | 20 |
| GO:MF | sodium channel activity | GO:0005272 | 0.000329408 | 40 | 7 |
| GO:MF | potassium ion transmembrane transporter activity | GO:0015079 | 0.00037568 | 153 | 12 |
| GO:MF | channel activity | GO:0015267 | 0.000598538 | 473 | 21 |
| GO:MF | passive transmembrane transporter activity | GO:0022803 | 0.000598538 | 473 | 21 |
| GO:MF | gated channel activity | GO:0022836 | 0.000659292 | 324 | 17 |
| GO:MF | extracellular matrix structural constituent conferring tensile strength | GO:0030020 | 0.000687988 | 35 | 6 |
| GO:MF | inorganic cation transmembrane transporter activity | GO:0022890 | 0.000711873 | 560 | 23 |
| GO:MF | signaling receptor binding | GO:0005102 | 0.00107475 | 1667 | 22 |
| GO:MF | ion binding | GO:0043167 | 0.003032465 | 5890 | 98 |
| GO:MF | extracellular matrix structural constituent | GO:0005201 | 0.00484187 | 136 | 10 |
| GO:MF | protein binding | GO:0005515 | 0.005151378 | 10433 | 146 |
| GO:MF | transmembrane transporter activity | GO:0022857 | 0.006105685 | 1070 | 32 |
| GO:MF | voltage-gated cation channel activity | GO:0022843 | 0.008534612 | 140 | 4 |
| GO:MF | cation channel activity | GO:0005261 | 0.010938433 | 322 | 15 |
| GO:MF | enzyme binding | GO:0019899 | 0.012676422 | 2254 | 6 |
| GO:MF | voltage-gated ion channel activity | GO:0005244 | 0.017886223 | 189 | 11 |
| GO:MF | voltage-gated channel activity | GO:0022832 | 0.018765674 | 190 | 11 |
| GO:MF | transporter activity | GO:0005215 | 0.029185018 | 1161 | 32 |
| GO:MF | sterol response element binding | GO:0032810 | 0.030131506 | 3 | 2 |

**Table S4. Cytokine/Chemokine 44-Plex Discovery Assay® Array of Serum of *Mecp2 Dup* and wild type mice the day prior (d.p.i -1) to *Influenza* infection.** Statistical analysis performed with Student's t-test.

**Chemokines and cytokines analysis, Serum d.p.i. -1**

| **Molecule** | **Concentration (pg/mL +/- SEM)** | | **Fold Change** | **p value** |
|---|---|---|---|---|
| | **Wild type** | ***Mecp2 Dup*** | | |
| Eotaxin | 574.56 ± 71.49 | 530.48 ± 57.85 | 0.92 | 0.6404 |
| G-CSF | 280.67 ± 54.98 | 422.94 ± 85.33 | 1.51 | 0.1864 |
| GM-CSF | 14.64 ± 2.64 | 11.65 ± 4.33 | 0.80 | 0.5453 |
| IFNγ | 157.61 ± 154.7 | 39.89 ± 20.56 | 0.25 | 0.4652 |
| IL-1α | 526.09 ± 58.09 | 431.11 ± 45.85 | 0.82 | 0.2236 |
| IL-1β | 3.75 ± 0.85 | 5.25 ± 1.06 | 1.40 | 0.2926 |
| IL-2 | 4.72 ± 0.45 | 4.05 ± 0.55 | 0.86 | 0.3674 |
| IL-3 | 2.39 ± 0.8 | 1.75 ± 0.64 | 0.73 | 0.5592 |
| IL-4 | 0.14 ± 0.06 | 0.26 ± 0.05 | 1.88 | 0.1972 |
| IL-5 | 3.59 ± 0.67 | 9.92 ± 5.55 | 2.76 | 0.2804 |
| IL-6 | 5.73 ± 2.52 | 8.17 ± 1.63 | 1.43 | 0.4326 |
| IL-7 | 2.78 ± 0.5 | 2.17 ± 0.52 | 0.78 | 0.4148 |
| IL-9 | 22.86 ± 2.45 | 20.56 ± 2.37 | 0.90 | 0.5140 |
| IL-10 | 10.94 ± 1.08 | 8.85 ± 1.02 | 0.81 | 0.1850 |
| IL-12p40 | 17.85 ± 3.48 | 9.79 ± 1.82 | 0.55 | 0.0632 |
| IL-12p70 | 17.71 ± 4.25 | 16.05 ± 4.33 | 0.91 | 0.7914 |
| IL-13 | 100.01 ± 32.9 | 63.93 ± 4.12 | 0.64 | 0.2980 |
| IL-15 | 79.48 ± 11.44 | 56.91 ± 11.8 | 0.72 | 0.1950 |
| IL-17 | 2.45 ± 0.36 | 2.04 ± 0.26 | 0.83 | 0.3827 |
| CXCL10 | 74.47 ± 9.8 | 74.55 ± 8.47 | 1.00 | 0.9953 |
| CXCL1 | 55.23 ± 9.39 | 55.21 ± 2.94 | 1.00 | 0.9984 |
| LIF | 0.34 ± 0 | Not detected | N/A | N/A |
| CXCL5 | 3225.31 ± 279.55 | 3110.24 ± 240.17 | 0.96 | 0.7602 |
| CCL2 | 25.78 ± 2.84 | 26.96 ± 1.8 | 1.05 | 0.7338 |
| M-CSF | 20.07 ± 4.92 | 139.09 ± 100.7 | 6.93 | 0.2607 |
| CXCL9 | 1005.04 ± 338.93 | 1262.18 ± 141.78 | 1.26 | 0.4973 |
| CCL3 | 59.49 ± 7.48 | 54.54 ± 10.32 | 0.92 | 0.7048 |
| CCL4 | 109.57 ± 11.76 | 106.91 ± 6.59 | 0.98 | 0.8473 |
| CXCL2 | 709.09 ± 1.33 | 707.98 ± 1.21 | 1.00 | 0.5511 |
| CCL5 | 35.31 ± 3.05 | 43.53 ± 2.84 | 1.23 | 0.0725 |
| TNFα | 14 ± 0.75 | 13.53 ± 0.59 | 0.97 | 0.6295 |
| VEGF | 1.65 ± 0.27 | 1.41 ± 0.08 | 0.85 | 0.4096 |

| | | | | |
|---|---|---|---|---|
| CCL21 | 17540.11 ± 659.84 | 17145.67 ± 930.58 | 0.98 | 0.7355 |
| EPO | 1011.68 ± 212.34 | 678.58 ± 82.56 | 0.67 | 0.1694 |
| CX3CL1 | 616.49 ± 53.76 | 455.24 ± 25.97 | 0.74 | 0.0193 |
| IFNβ-1 | 1703.83 ± 388.09 | 1680.41 ± 672.22 | 0.99 | 0.9764 |
| IL-11 | 44.73 ± 9.71 | 30.45 ± 2.21 | 0.68 | 0.1771 |
| IL-16 | 5874.93 ± 949.12 | 5103.9 ± 723.72 | 0.87 | 0.5304 |
| IL-20 | 1801.93 ± 297.37 | 2090.13 ± 474.65 | 1.16 | 0.6162 |
| CCL12 | 337.97 ± 77.18 | 440.08 ± 70.03 | 1.30 | 0.3466 |
| CCL22 | 514.89 ± 136.58 | 566.6 ± 84.56 | 1.10 | 0.7531 |
| CCL20 | 56.07 ± 6.69 | 52.29 ± 8.39 | 0.93 | 0.7313 |
| CCL19 | 808.81 ± 92.72 | 447.21 ± 89.67 | 0.55 | 0.0159 |
| CCL17 | 116.51 ± 25.59 | 104.08 ± 25.08 | 0.89 | 0.7348 |
| TIMP-1 | 5405.51 ± 577.66 | 6651.72 ± 516.47 | 1.23 | 0.1338 |

**Table S5. Cytokine/Chemokine 44-Plex Discovery Assay® Array of Serum of *Mecp2 Dup* and wild type mice at day 4 post *Influenza* infection (d.p.i +4).** Statistical analysis performed with Student's t-test.

**Chemokines and cytokines analysis, Serum d.p.i. +4**

| **Molecule** | **Concentration (pg/mL +/- SEM)** | | **Fold Change** | **p value** |
|---|---|---|---|---|
| | **Wild type** | ***Mecp2 Dup*** | | |
| Eotaxin | 1803.42 ± 137.49 | 1801.73 ± 166.79 | 0.9991 | 0.9939 |
| G-CSF | 597.17 ± 155.25 | 628.73 ± 80.66 | 1.0529 | 0.8599 |
| GM-CSF | 10.57 ± 1.42 | 19.94 ± 4.28 | 1.8860 | 0.0303 |
| IFNγ | 51.01 ± 20.39 | 113.69 ± 55.27 | 2.2288 | 0.3083 |
| IL-1α | 310.14 ± 29.26 | 319.15 ± 53.69 | 1.0290 | 0.8853 |
| IL-1β | 0.91 ± 0.18 | 1.4 ± 0.73 | 1.5245 | 0.4135 |
| IL-2 | 4.65 ± 0.43 | 3.78 ± 0.64 | 0.8125 | 0.2840 |
| IL-3 | 1.69 ± 0.73 | 4.31 ± 1.78 | 2.5499 | 0.1774 |
| IL-4 | 0.22 ± 0.15 | 0.22 ± 0.1 | 1.0000 | 1.0000 |
| IL-5 | 7.28 ± 2.64 | 3.59 ± 0.52 | 0.4940 | 0.1964 |
| IL-6 | 103.27 ± 37.84 | 118.4 ± 28.65 | 1.1465 | 0.7554 |
| IL-7 | 2.06 ± 0.74 | 2.84 ± 0.73 | 1.3767 | 0.4895 |
| IL-9 | 21.72 ± 2.28 | 24.04 ± 1.27 | 1.1068 | 0.3930 |
| IL-10 | 7.67 ± 0.78 | 8.67 ± 0.85 | 1.1302 | 0.4061 |
| IL-12p40 | 12.83 ± 2.01 | 16.21 ± 2.91 | 1.2632 | 0.3586 |
| IL-12p70 | 17.44 ± 7.16 | 29.25 ± 13.27 | 1.6768 | 0.4803 |
| IL-13 | 136.7 ± 35.95 | 129.92 ± 10.54 | 0.9504 | 0.8594 |
| IL-15 | 82.48 ± 12.29 | 78.63 ± 15.61 | 0.9533 | 0.8496 |
| IL-17 | 1.56 ± 0.34 | 1.74 ± 0.55 | 1.1160 | 0.7859 |
| CXCL10 | 428.84 ± 47.29 | 406.07 ± 42.03 | 0.9469 | 0.7252 |
| CXCL1 | 93.13 ± 9.93 | 115.82 ± 18.95 | 1.2436 | 0.3099 |
| LIF | 1.01 ± 0 | 0.73 ± 0 | 0.7228 | N/A |
| CXCL5 | 4302.98 ± 347.89 | 5165.44 ± 631.54 | 1.2004 | 0.2547 |
| CCL2 | 55.77 ± 9.19 | 65.39 ± 14.27 | 1.1724 | 0.5816 |
| M-CSF | 20.06 ± 4.88 | 36.2 ± 13.43 | 1.8046 | 0.2810 |
| CXCL9 | 2119.18 ± 266.85 | 3523.4 ± 330.05 | 1.6626 | 0.0062 |
| CCL3 | 136.99 ± 8.58 | 108.38 ± 14.55 | 0.7912 | 0.1162 |
| CCL4 | 165.12 ± 9.44 | 170.38 ± 11.23 | 1.0318 | 0.7264 |
| CXCL2 | 707.25 ± 0.83 | 707.62 ± 0.99 | 1.0005 | 0.7811 |
| CCL5 | 39.73 ± 1.24 | 56.05 ± 4.07 | 1.4109 | 0.0024 |
| TNFα | 18.94 ± 1.02 | 20.26 ± 1.78 | 1.0698 | 0.5322 |
| VEGF | 1.74 ± 0.22 | 1.91 ± 0.12 | 1.0982 | 0.5264 |

| | | | | |
|---|---|---|---|---|
| CCL21 | 22250.88 ± 343.44 | 20570.85 ± 831.44 | 0.9245 | 0.0864 |
| EPO | 924.65 ± 186.86 | 722.85 ± 105.92 | 0.7818 | 0.3660 |
| CX3CL1 | 907.75 ± 87.78 | 795.34 ± 146.35 | 0.8762 | 0.5225 |
| IFNβ-1 | 1960.96 ± 809.53 | 2560.38 ± 1003.13 | 1.3057 | 0.6502 |
| IL-11 | 84 ± 19.59 | 50.27 ± 6.81 | 0.5984 | 0.1299 |
| IL-16 | 9859.11 ± 625.11 | 7511.15 ± 725.21 | 0.7618 | 0.0305 |
| IL-20 | 1482.15 ± 154.04 | 2290.45 ± 333.31 | 1.5454 | 0.0480 |
| CCL12 | 1446.66 ± 145.53 | 1535.04 ± 217.89 | 1.0611 | 0.7417 |
| CCL22 | 1372.61 ± 332.07 | 1518.94 ± 174.7 | 1.1066 | 0.7034 |
| CCL20 | 222.74 ± 51.15 | 203.44 ± 55.1 | 0.9133 | 0.8017 |
| CCL19 | 906.41 ± 183.14 | 940.89 ± 135.61 | 1.0380 | 0.8823 |
| CCL17 | 369.97 ± 61.43 | 320.12 ± 35.28 | 0.8653 | 0.4951 |
| TIMP-1 | 15313.99 ± 638.45 | 14257.63 ± 506.22 | 0.9310 | 0.2192 |

**Table S6. Cytokine/Chemokine 44-Plex Discovery Assay® Array of BALF of *Mecp2 Dup* and wild type mice at day 4 post *Influenza* infection (d.p.i +4).** Statistical analysis performed with Student's t-test.

**Chemokines and cytokines analysis, BALF d.p.i. +4**

| Molecule | Concentration (pg/mL +/- SEM) | | Fold Change | p value |
|---|---|---|---|---|
| | **Wild type** | ***Mecp2 Dup*** | | |
| Eotaxin | 150.65 ± 10.22 | 119.88 ± 4.1 | 0.80 | 0.0162 |
| G-CSF | 38.11 ± 6.08 | 55.03 ± 8.58 | 1.44 | 0.1339 |
| GM-CSF | 3.47 ± 0.95 | 4.85 ± 0.83 | 1.40 | 0.3045 |
| IFNγ | 352.07 ± 196.13 | 2204.69 ± 737.62 | 6.26 | 0.0319 |
| IL-1α | 3.2 ± 0.59 | 11.48 ± 3.26 | 3.59 | 0.0951 |
| IL-1β | 0.55 ± 0.19 | 3.24 ± 1.41 | 5.90 | 0.1667 |
| IL-2 | 0.53 ± 0.08 | 0.89 ± 0.06 | 1.69 | 0.0062 |
| IL-3 | 0.78 ± 0.05 | 0.84 ± 0.05 | 1.08 | 0.4309 |
| IL-4 | 0.05 ± 0 | 0.08 ± 0 | 1.57 | 0.0305 |
| IL-5 | 0.49 ± 0.11 | 0.6 ± 0.04 | 1.22 | 0.4006 |
| IL-6 | 253.66 ± 65.72 | 242.48 ± 57.93 | 0.96 | 0.9006 |
| IL-7 | 0.95 ± 0.22 | 1.17 ± 0.18 | 1.23 | 0.4648 |
| IL-9 | 3.51 ± 0.45 | 4.41 ± 0.9 | 1.25 | 0.3940 |
| IL-10 | 0.28 ± 0.07 | 0.94 ± 0.6 | 3.31 | 0.1945 |
| IL-12p40 | Not detected | Not detected | N/A | N/A |
| IL-12p70 | 1.44 ± 0.35 | 1.78 ± 0.4 | 1.24 | 0.5473 |
| IL-13 | 0.8 ± 0.12 | 1.94 ± 0.47 | 2.43 | 0.0427 |
| IL-15 | 2.31 ± 0.81 | 3.09 ± 0.75 | 1.34 | 0.5033 |
| IL-17 | 0.15 ± 0.02 | 0.36 ± 0.08 | 2.33 | 0.0551 |
| CXCL10 | 515.73 ± 96.37 | 991.88 ± 194.33 | 1.92 | 0.0486 |
| CXCL1 | 93.62 ± 17.76 | 114.84 ± 10.4 | 1.23 | 0.3231 |
| LIF | 6.24 ± 1.83 | 5.86 ± 0.85 | 0.94 | 0.8564 |
| CXCL5 | Not detected | Not detected | N/A | N/A |
| CCL2 | 95.2 ± 18.5 | 77.81 ± 6.49 | 0.82 | 0.3929 |
| M-CSF | 1.32 ± 0.23 | 1.95 ± 0.1 | 1.47 | 0.0331 |
| CXCL9 | 72.41 ± 16.8 | 230.72 ± 47.81 | 3.19 | 0.0088 |
| CCL3 | 93.9 ± 16.19 | 112 ± 7.19 | 1.19 | 0.3272 |
| CCL4 | 134.58 ± 22.71 | 145.71 ± 16.74 | 1.08 | 0.7004 |
| CXCL2 | 14.98 ± 2.29 | 27.47 ± 2.26 | 1.83 | 0.0022 |
| CCL5 | 1.91 ± 0.41 | 3.29 ± 0.68 | 1.72 | 0.1109 |
| TNFα | 3.75 ± 0.68 | 8.19 ± 1.65 | 2.19 | 0.0291 |
| VEGF | 4.44 ± 1.37 | 6.43 ± 1.51 | 1.45 | 0.3494 |

| | | | | |
|---|---|---|---|---|
| CCL21 | 8255.63 ± 3702.34 | 6379.5 ± 2506.59 | 0.77 | 0.6715 |
| EPO | Not detected | Not detected | N/A | N/A |
| CX3CL1 | 17.15 ± 2.44 | 20.7 ± 1.97 | 1.21 | 0.2808 |
| IFNβ-1 | 60.27 ± 5.32 | 74.43 ± 6.65 | 1.23 | 0.1226 |
| IL-11 | 0 ± 0 | 0 ± 0 | N/A | N/A |
| IL-16 | 253.77 ± 37.57 | 273.49 ± 15.13 | 1.08 | 0.6353 |
| IL-20 | 27.83 ± 2.18 | 51.78 ± 2.48 | 1.86 | 0.0006 |
| CCL12 | 231 ± 27.72 | 252.59 ± 14.07 | 1.09 | 0.5007 |
| CCL22 | 46.05 ± 7.14 | 63.24 ± 11.69 | 1.37 | 0.2336 |
| CCL20 | 2.48 ± 0.97 | 3.28 ± 0.94 | 1.32 | 0.5660 |
| CCL19 | 34.89 ± 3.71 | 55.75 ± 3.58 | 1.60 | 0.0016 |
| CCL17 | 12.47 ± 1.74 | 11.83 ± 1.48 | 0.95 | 0.7830 |
| TIMP-1 | 2822.88 ± 493.37 | 1903.41 ± 68.44 | 0.67 | 0.0897 |

**Table S7. sgRNAs and bridging donor utilized to generate the *Mecp2 Dup* mice**

| **Sequence name** | **Sequence** |
|---|---|
| Bridging donor | GTTGCTCCCACCTGAACAGCAAAGTAGAGTTGGTCTTGGTTGC<br>AGAGGGCAGGGGTGGATGAAGGTGAGTCAGCTCTGAGAGTTA<br>GAGAGTGGGAAAGCTGGCCCAGCCCCTTGCTTGCTGCAGGAA<br>ACTCATCTGgaattcTAGTCGGCAGAGGTGGCAAGAAACTGAAG<br>TAAGGGCTGGTGAGATGGCTCAGCGGTTAAAGAGCGCCGACT<br>GCTCTTCCAAAGGTCATGAGTTCAAATCCCAGCAACCACATG<br>GTGGCTCATAACCATCTGTAATGAAATCTGATGCCCTCTTCTG<br>GTGTGTCTGAAGACAGCTACAGTGTACTTACAT |
| Irak1_sgRNA | TAGCATCAATCAGCCCTAGT |
| Tex28_sgRNA | CAGCTGTACTATGTTACCCAG |

**Table S8. Oligonucleotides utilized in this study**

| Primer name | Sequence | Purpose |
|---|---|---|
| mMecp2 Dup 5' del F | AACTCACCAGAGGTAGTGGAAAAG | 5' breakpoint PCR + sequencing |
| mMecp2 Dup 5' del R | CTATAGCCCAGGCTTACAGTGATT | |
| mMecp2 geno F | ATGGTTGGAGCCTGGGTATT | Dup junction PCR + sequencing |
| mMecp2 geno R | CCATCTCACCAGCCCTTACT | |
| mMecp2 3' break | ATGGTGAGATGACCAATGACTTC | 3' breakpoint PCR + sequencing |
| mMecp2 Dup 3' del R | CCTGGAGATCTTGTAGTTTTGGAT | |
| mMecp2 Del F | GAGTAACTCCTGTCTGTGTTGTCT GG | Del junction PCR + sequencing |
| mMecp2 Del R | CTTACATTGCAGAGTTGAGCTGCGTT | |
| mMecp2 e1 F | AGG AGA GAC TGG AGG AAA AGT C | RT-PCR *Mecp2* isoforms |
| mMecp2 e2 F | CTTAAACTTCAGTGGCTTGTCTCTG | |
| mMecp2 e1-e2 R | CTCACCAGTTCCTGCTTTGATGT | |
| mMecp2 qPCR ex3 F2 | CGATCTGCTGGAAAGTATGATGT | qPCR |
| mMecp2 qPCR ex4 R2 | CTTCTTAGGTGGTTTCTGCTCTCT | |
| mIrak1 qPCR F | TGTGAAGAGACTGAAGGAGGAAG | |
| mIrak1 qPCR R | CCGCAAAGTCTACGATATTTGG | |
| mGapdh F | TGTTTGTGATGGGTGTGAACC | |
| mGapdh R | ACTGTGGTCATGAGCCCTTC | |
| mCacna1g qPCR F | GACCATGTGGTCCTCGTCATCA | |
| mCacna1g qPCR R | TTTCAGCCAGGAAGACTGCCGT | |
| mGad2 qPCR F | CATTGATAAGTGTTTGGAGCTAGCA | |
| mGad2 qPCR R | GTGCGCAAACTAGGAGGTACAA | |
| mFxyd7 qPCR F | AAGGCGGATTCCAGGTCTGA | |
| mFxyd7 qPCR R | GGAGGGCAGTTCCGACTTAC | |
| mPcbp3 qPCR F | GACGCCATATTCCAGTGTGTG | |
| mPcpb3 qPCR R | GTCTGTCCGAGGGGAGGAAA | |
